# Human-engineered heart tissues recapitulate tissue-scale mechanisms underlying ventricular tachycardia

**DOI:** 10.64898/2026.02.11.705466

**Authors:** Matthew Fiedler, Alejandra Vasquez Limeta, Ernesto Reyes Sanchez, Leah Carter, Francisco Altamirano

## Abstract

Human iPSC-derived engineered heart tissues (EHTs) and cardiac organoids are increasingly used to model cardiac physiology and drug responses, yet it remains unclear whether they can reproduce tissue-scale mechanisms underlying ventricular arrhythmias. Advanced electrophysiological characterization of EHTs has been limited by the lack of a recording framework compatible with small, perfused preparations, despite the availability of mapping hardware.

We present a reproducible workflow that couples milliPillar-based EHT fabrication with high-resolution (22 µm spatial, 1 ms temporal) dual channel optical mapping using RH-237 for voltage and Rhod-2 AM for Ca²⁺. Baseline electrophysiological measurements align with published data from human and animal cardiac tissues, showing rate-dependent restitution of action potential and Ca²⁺ transient duration (APD and CaD), conduction slowing at higher pacing rates, and physiologic AP-Ca²⁺ activation latency. Selective hERG blockade with E-4031 prolongs APD, confirming pharmacological sensitivity.

To interrogate mechanisms underlying ventricular tachycardia (VT), we utilized an established proarrhythmic perturbation widely validated in animal models of acquired long-QT syndrome, combining hERG inhibition with hypokalemia and hypomagnesemia; electrolyte disturbances commonly encountered in clinical settings. Treated EHTs (VT group) displayed tachyarrhythmic contractile bursts with marked beat-to-beat instability, whereas controls responded synchronously to field stimulation. Beat-resolved optical mapping revealed progressive diastolic interval shortening, APD dispersion with transient localized long-short APD zones, and regional depression of excitability that together formed spatial conduction barriers precipitating wavebreak and reentry. Early afterdepolarizations contributed to triggered activity and created localized repolarization barriers at long-short APD zones, leading to rotor formation. Phase singularity tracking identified short-lived rotors localized predominantly in the heads of VT EHTs and absent in controls. A minority of tissues exhibited multiple simultaneous rotors and wavelets generating chaotic-like activation. Although tissue acceleration promoted rotor formation, these events were brief, likely due to the spatial limitations of EHTs, and treated tissues more closely resembled VT than Torsades de Pointes or sustained fibrillation.

Our comprehensive studies demonstrate that human iPSC-derived EHTs can recapitulate the tissue-scale VT mechanisms associated with acquired long QT syndrome – APD dispersion with long-short APD zones, triggered activity, conduction block, wavebreak and reentry – which had previously been assessed only in intact hearts. The presented platform thus provides a scalable, non-animal system for mechanistic arrhythmia research.

## 1. Introduction

Ventricular tachycardia (VT) and its degeneration into ventricular fibrillation (VF) remain the primary drivers of sudden cardiac death.^1^ The clinical presentation is diverse including genetic disorders such as long QT syndrome (LQTS) and as a bidirectional interaction with heart failure, where VT leads to heart dysfunction and pathological remodeling that further increases cardiovascular mortality.^2,3^ At the cellular level, VT arises from multiple electrophysiological alterations including rate-dependent shortening of the diastolic interval (DI), action potential duration (APD) spatial and temporal heterogeneity, slowing of conduction velocity (CV), conduction block, wavebreak and the formation of re-entrant circuits.^4^ Early and delayed afterdepolarizations (EADs and DADs) can further aggravate these mechanisms by generating triggered activity which results in premature beats.^5,6^ High-resolution optical mapping of voltage or Ca²⁺ signals in animal hearts and human preparations from donor hearts has been indispensable for dissecting these electrophysiological spatiotemporal dynamics and elucidating arrhythmia mechanism.^7^ However, species-specific baseline electrophysiological differences^8^, intrinsic human genetic diversity and human cardiac remodeling during disease can promote divergent mechanistic paths, which has motivated human-based platforms to bridge the translation gap.^9–11^

Human induced pluripotent stem cell-derived cardiomyocytes (hiPSC-CMs) provide scalable access to patient-specific electrophysiology and have revolutionized disease modelling and drug-cardiotoxicity screening.^12,13^ Yet most hiPSC-CM electrophysiological studies are confined to single-cell recordings, monolayers, or organoid assays that rely on measurements which, although informative for cellular mechanisms, lack spatiotemporal resolution.^12^ Moreover, these reductionistic platforms lack the structural anisotropy, mechanical load, and multicellular connectivity required to initiate and sustain the tissue-scale phenomena that underlie sustained VT/VF.

Engineered heart tissues (EHTs) that embed hiPSC-CMs and fibroblasts in a three-dimensional format represent an intermediate step between reductionist models and intact hearts. By recapitulating mechanical loading and multicellular composition, EHTs display more physiological excitation-contraction coupling (ECC) and tissue properties.^14–16^ Nevertheless, high spatiotemporal resolution electrophysiological interrogation of human EHTs remains limited for two principal reasons: (i) the lack of a reproducible, high-throughput fabrication pipeline that yields tissue constructs compatible with optical mapping studies; and (ii) the difficulty of establishing stable imaging conditions including continuous perfusion, precise temperature control, minimized motion artifacts and synchronized electrical stimulation required for quantitative voltage and Ca²⁺ mapping at millisecond temporal resolution. There is still a gap in knowledge as to whether hiPSC-CM-based EHTs can faithfully reproduce the substrate, initiators, and drivers of human arrhythmias leading to VT/VF, or whether they are limited to artifactual and/or simplified versions of in vitro rhythmic disturbances.

To address this gap, we established a robust workflow for high-resolution, simultaneous optical mapping of voltage and Ca²⁺ in EHTs. By incorporating modifications to the open-source milliPillar EHT platform^17^ we facilitated mold generation for PDMS casting and improved plating conditions for dual-voltage and Ca²⁺ optical mapping. We developed a compatible 3D-printed imaging stage that provides perfusion, temperature control, and electrical stimulation while minimizing motion artifacts in buoyant PDMS-based EHT constructs, using a commercially available optical mapping recording system. Using this system, we first characterized baseline electrophysiological responses in EHTs at an unprecedented spatiotemporal resolution (∼22 µm spatial and 1 ms temporal) and observed results similar to those from previously published adult hearts. We then challenged the tissues with a pro-arrhythmic milieu: low extracellular K⁺/Mg²⁺ concentration (low K⁺/Mg²⁺) combined with selective hERG blockade, which reproduces acquired LQTS complications leading to VT and the highly lethal arrhythmia Torsades de Pointes (TdP).^18,19^

Analysis of beat-to-beat APD, DI, and upstroke velocity together with phase-singularity and wavelet tracking reveals that the EHTs develop restitution-driven recovery failure, localized conduction slowing, functional block, and wavebreak, culminating in short-lived re-entrant rotors. These alterations were driven by localized and transient APD dispersion gradients that created long-short APD regions, a known substrate for initiating human arrhythmias. These observations demonstrate that hiPSC-CM EHTs can faithfully recapitulate the multidimensional substrate of VT with limitations in VF modeling.

## 2. Results

### 2.1. Implementation of modified individual milliPillar modules

To increase throughput and compatibility with standard cell culture workflows, minor modifications were made to the milliPillar platform.^17^ Comprehensive details of mold modifications, print orientation, post-processing, and STL file are in the Supplemental data (**Suppl. Table 1-2**, **STL File 1**, and **Suppl. Fig. 1**). These changes allow multiple mold fabrication to be printed on a low-cost consumer resin printer (**Fig. 1A**). After PDMS casting, individual milliPillar modules were trimmed and attached to multi-well plates, facilitating plating, media exchange, and detachable intact EHTs for optical mapping (**Fig. 1B, Suppl. Fig. 2**, and **Video 1**). This modularity enabled rapid iteration of handling and imaging configurations (e.g., longitudinal electrical pacing with C-Pace, IonOptix followed by terminal optical mapping) without affecting tissue structure or function.

**Figure 1.**
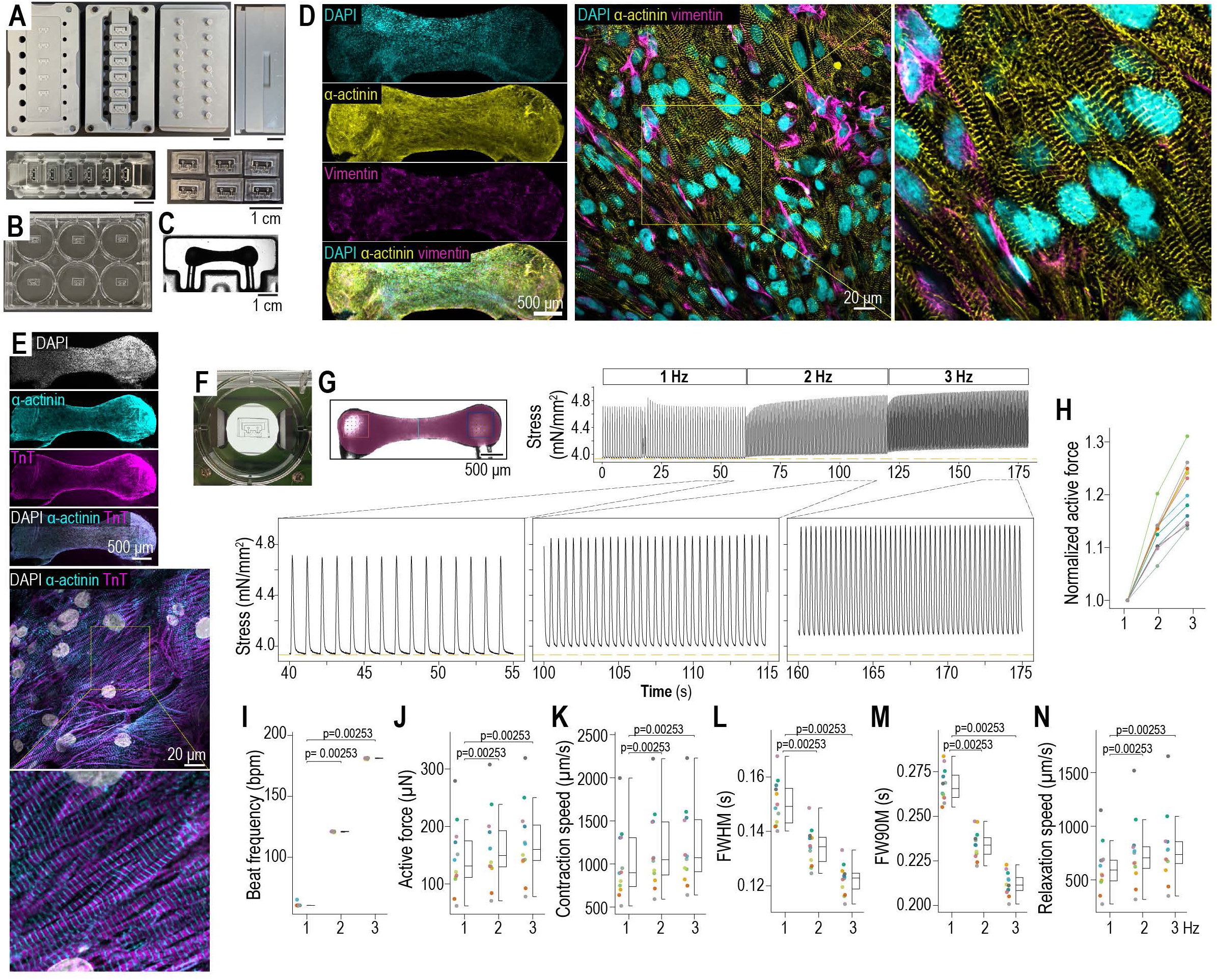
Reproducible fabrication of EHTs using a modified milliPillar platform. **A**. Modified milliPillar molds were 3D-printed in temperature-resistant resin and used to cast PDMS. **B**. Individual modules were attached to multi-well dishes, increasing flexibility and compatibility with standard electrophysiology and cell culture workflows. **C**. Representative brightfield image of an EHT 1 month after plating. **D**. Confocal section stained for DAPI, α-actinin, and vimentin; magnified insert shows longitudinal cardiomyocyte alignment with interspersed fibroblasts. **E**. Confocal section stained for DAPI, α-actinin, and troponin T (TnT). Magnified insert reveals adult-like contractile filament organization. **F**. IonOptix C-Pace insert was used to deliver field electrical stimulation. **G**. Contractility was assessed from 200-fps brightfield videos under stimulation at 37 °C in maturation media; the final 15 s of each pacing frequency plus a 5 s padding were analyzed with BeatProfiler. An increase in the force-frequency relationship was observed in all EHTs (**H**). **I**-**N**. Quantifications are shown as box plots (median, interquartile range (IQR), and 1.5X IQR whiskers) overlaid with scatter points for N = 12 independent EHTs from 2 batches, paired by frequency. Several endpoints were non-normally distributed (Shapiro-Wilk test), so Friedman test followed by paired Wilcoxon test versus the 1 Hz condition were applied for **I-N**. Repeated p-values reflect identical signed-rank patterns across outcomes.

EHTs were fabricated by mixing hiPSC-CMs and human cardiac fibroblasts at a 3:1 ratio in collagen hydrogel. Using the original protocol^1^, we initially detached tissues with a 26-gauge needle at 1 and 24 h post-plating; in our hands, however, needle scoring slowed plating, introduced user- and batch-dependent variability, increased assembly failures, and left PDMS strands in the tissue. To eliminate manual scoring, we applied corona plasma treatment followed by KOSR blocking to reduce PDMS-hydrogel adhesion (**Suppl. Table 3**). With this conditioning, 100% of EHTs self-detached spontaneously within 24 h post-plating (**Suppl. Fig. 2F**). We then transitioned to DMEM-based metabolic maturation media supplemented with KOSR as a lipid source, with additional supplements as described.^20^

After transition to metabolic maturation media, self-assembly began within 24 h and spontaneous contractions appear after ∼72 h. One-week EHT formation was 79 ± 9.3% (four batches) with failures occurring when tissues could not maintain tension on one pillar head, collapsing into spheroids on the other. To improve electrophysiological responses, hormonal maturation with triiodothyronine (T3) and dexamethasone was applied for 2 weeks^21^ starting at week 2, after which EHTs returned to DMEM-based metabolic maturation media until experimentation.

To initially validate EHT organization and function, we performed immunostaining and contractility measurements in 1 month-old EHTs. Immunostaining confirmed longitudinal cardiomyocyte alignment with interspersed, elongated fibroblasts, recapitulating the anisotropic structure of mature adult myocardium (**Fig. 1D-E**). For contractility assays, EHTs were paced in culture media at 1, 2, and 3 Hz for 60 sec intervals using an IonOptix C-Pace with brightfield recordings at 200 frames per second (**Fig. 1F, Suppl. Video 2**). Contractility and kinetics were assessed using BeatProfiler.^22^ EHTs exhibited a positive force-frequency relationship with consistent 1:1 capture and failure rate of <5% at 1 Hz stimulation (**Fig. 1H-I**). Active force and contraction/relaxation kinetics were comparable to previously published milliPillar data (**Fig. 1J-N**).^17,22,23^ These data indicate that EHTs are functionally mature by 1 month, prompting initial electrophysiological characterization at this time point.

### 2.2. Development of a reproducible, high-resolution, simultaneous voltage and Ca²⁺ optical mapping system

High spatiotemporal resolution optical mapping requires unobstructed optical access to the entire tissue preparation with motion artifacts minimization, while tissues are buoyant and require stable temperatures and near-constant perfusion to maintain viability. We thus developed a simple and adaptable imaging workflow for milliPillar-based EHT modules to address these requirements.

Dye loading was optimized to maximize signal-to-noise ratio while minimizing phototoxicity; the potentiometric dye RH-237 (30 μM) and Ca²⁺-sensitive dye Rhod-2AM (10 μM) were co-loaded with 0.1% Pluronic F-127 in blebbistatin-containing (5 μM) Tyrode’s Solution. Loading for 20 min at 37 °C yielded robust voltage and Ca²⁺ signals, with longer incubations increasing phototoxicity and rendering tissues unresponsive to electrical pacing.

An Arduino-controlled transistor-transistor logic (TTL) signal generator was used to control electrical stimulation, LED illumination, camera acquisition, and to pause perfusion immediately before and during recording to minimize motion artifacts. Simultaneous dual-channel optical mapping was performed using a SciMedia MiCAM03 system (**Fig. 2A-B**). To secure the tissue during imaging, we designed a 3D-printed perfusion stage to stabilize individual milliPillar modules, maintain a constant level of physiological buffer at a constant temperature, and provide electrical stimulation to the tissues. A centrally positioned divot with adjustable EHT holder(s) secure milliPillar modules, while stable temperatures are maintained by a dual-channel ThermoClamp (AutoMate Scientific) heating perfused buffer and the imaging chamber (**Fig. 2C**). Electrical stimulation was delivered via platinum wires mounted on a slidable holder, enabling point stimulation of the right EHT head (**Fig. 2D**, **STL Files 2-7**).

**Figure 2.**
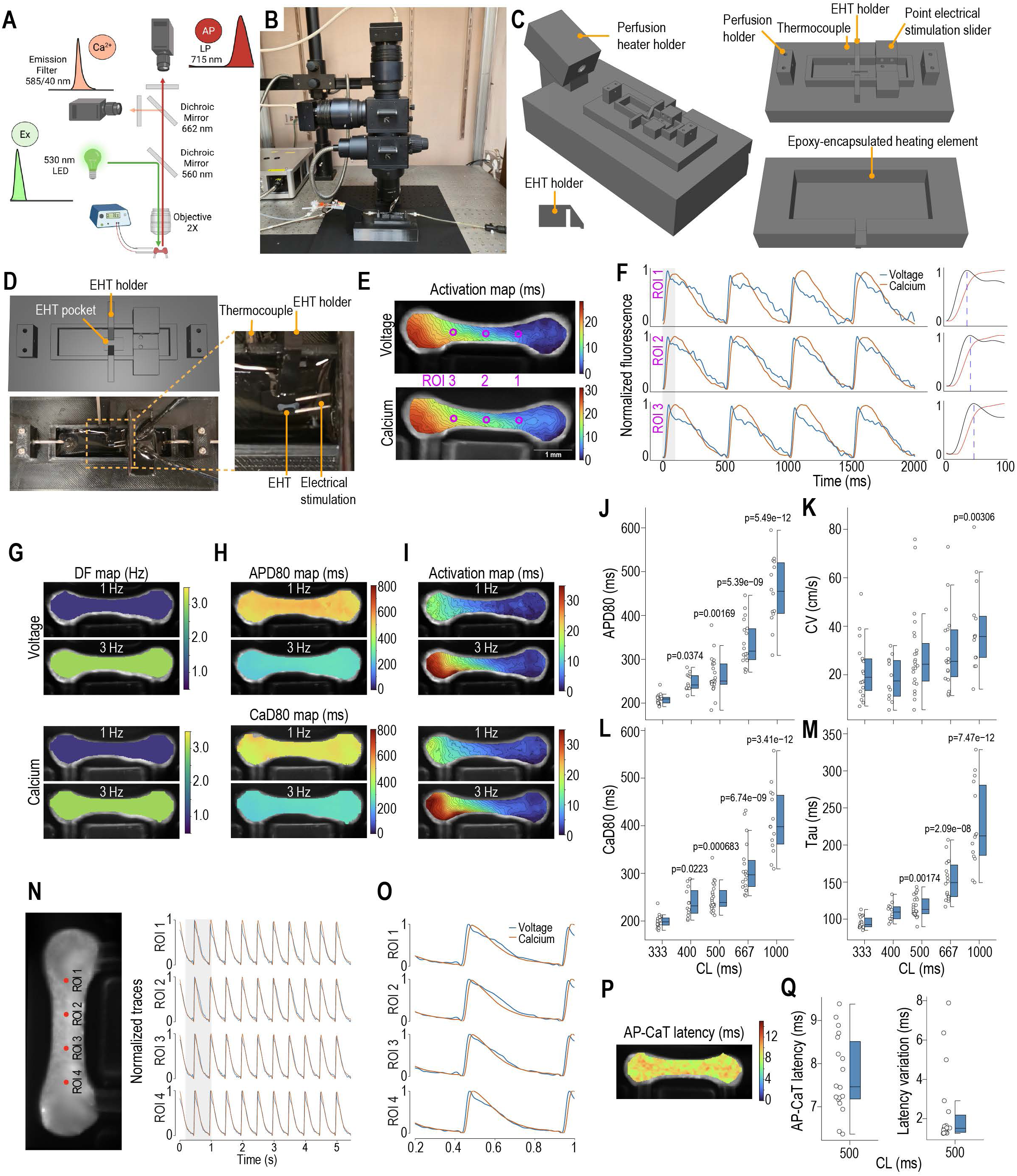
Optical mapping setup and EHT electrophysiology. **A**. Schematic representation. **B.** Photograph showing LED, mesoscope, camera and optics. **C**. 3D rendering of STL files for the imaging stage: a perfusion heater holder and epoxy-encapsulated heating elements maintain temperature during recording, while EHT holders and pocket minimize motion and flotation. An adjustable slider with two platinum wires provides point- or field-electrical stimulation. **D**. Top-down view showing milliPillar module positioning for a point-electrical stimulation experiment. **E**. Ensemble voltage (top) and Ca²⁺ (bottom) activation maps from a representative EHT paced at 2 Hz. Purple circles denote ROIs shown in panel **F,** showing normalized voltage (blue) and Ca²⁺ (orange) signals illustrating consistent propagation, with voltage preceding Ca²⁺. **G-I**. Dominant frequency (DF) and ensemble APD_80_, CaD_80_ and activation maps at 1 and 3 Hz. **J-M**. Quantifications of APD_80_/CV for voltage and CaD_80_/tau for Ca²⁺ signals reveal rate-dependent restitution curves. Data are expressed as cycle length (CL=(1/frequency) x 1000 ms). N = 11-18 EHTs from 4 batches. Several endpoints were non-normally distributed (Shapiro-Wilk), thus nonparametric testing was used for consistency. Because not all tissues were recorded at identical pacing frequencies (some experiments included 1-3 Hz with or without 0.5 intervals or had one excluded timepoint due to noise or artifacts), comparisons employed Kruskal-Wallis test with Dunn’s post hoc test versus the 333 ms condition. **N-O**. Voltage (blue) and Ca²⁺ (orange) traces from ROIs across a representative EHT paced at 2 Hz; the magnified region (0.2-1 s) on the right, shows AP-CaT latency across multiple ROIs. **P**. AP-CaT latency map generated with KairoSight shows uniform latency across the representative EHT from **N-O**. **Q**. Mean AP-CaT latency and variation (SD) indicate highly consistent ECC efficiency across tissues. Latency was calculated from all recordings presented in **J-M** obtained at 2 Hz (250 ms CL; N = 18 EHTs, 4 batches). All quantifications are shown as box plots (median, IQR, and 1.5X IQR whiskers) with overlaid scatter points.

Simultaneous voltage and Ca²⁺ recordings achieved 1 ms temporal resolution and ∼22 μm spatial resolution. Signal processing was achieved using spatial and temporal denoising, asymmetric least squares baseline correction, and 0-1 normalization (see Methods). EHTs exhibited uniform voltage and Ca²⁺ wave propagation from the right EHT head (point stimulation site) towards the left side, and a consistent action potential (AP) upstroke preceding the rise in Ca²⁺ transient (CaT) at every measured location (**Fig. 2E-F, Suppl. Video 3 and 4**).

### 2.3. Rate-dependent restitution curves and fast AP-Ca²⁺ coupling in EHTs

Voltage and Ca²⁺ signals revealed a homogeneous tissue response to 1-3 Hz pacing and expected physiological frequency-dependent changes (restitution curves) in action potential duration at 80% repolarization (APD_80_), head-to-head longitudinal conduction velocity (CV), Ca²⁺ duration at 80% baseline (CaD_80_), and CaT decay time constant (Tau); with all parameters decreasing as cycle length (CL=(1/frequency) x 1000) shortened, confirming rate-dependent electrophysiological adaptation (**Fig. 2G-M**). We also spatially evaluated excitation-contraction coupling (ECC) efficiency by measuring the latency between AP and CaT upstrokes (AP-CaT Latency) for the first beat of the recordings obtained at CL 500 ms (**Fig. 2N-O**). Mean AP-CaT latency of 7.7±0.2 ms were within ranges reported for adult human ventricular myocardium, supporting physiologically relevant ECC timings in EHTs. Collectively, these findings suggest that EHTs undergo electrophysiological maturation, with restitution curves and excitation-contraction coupling dynamics comparable to those of adult heart models. These points are further addressed in the Discussion section.

### 2.4. Pharmacological sensitivity to E4031

Pharmacological validation with the selective hERG blocker E4031 (100 nM) was performed at a baseline pacing 500 ms CL (2 Hz). Exposure to E4031 produced a reproducible prolongation of APD_80_ (**Suppl. Fig. 3**). Ensemble activation maps showed modest, non-significant elongation of activation time after drug treatment. Representative voltage traces from individual regions of interest illustrate the clear AP prolongation induced by E4031.

### 2.5. VT modeling in EHTs

VT can be initiated by triggered activity arising from repolarization abnormalities, as seen in acquired LQTS, which is associated with increased risk of life-threatening ventricular arrhythmias and sudden cardiac death.^24^ Combined high-dose hERG inhibition (500 nM) with hypokalemia and hypomagnesemia is an established pro-arrhythmic intervention that reliably induces VT-relevant phenomena.^18^ We used this model to determine whether EHT reproduce these clinically and mechanistically relevant arrhythmogenic behaviors by measuring contractility and electrophysiology (**Fig. 3A**). A modified Tyrode’s solution containing 2 mM K⁺ and 0.5 mM Mg²⁺ (reductions of 60% and 50%, respectively) supplemented with 500 nM E-4031, was used to induce VT.

**Figure 3.**
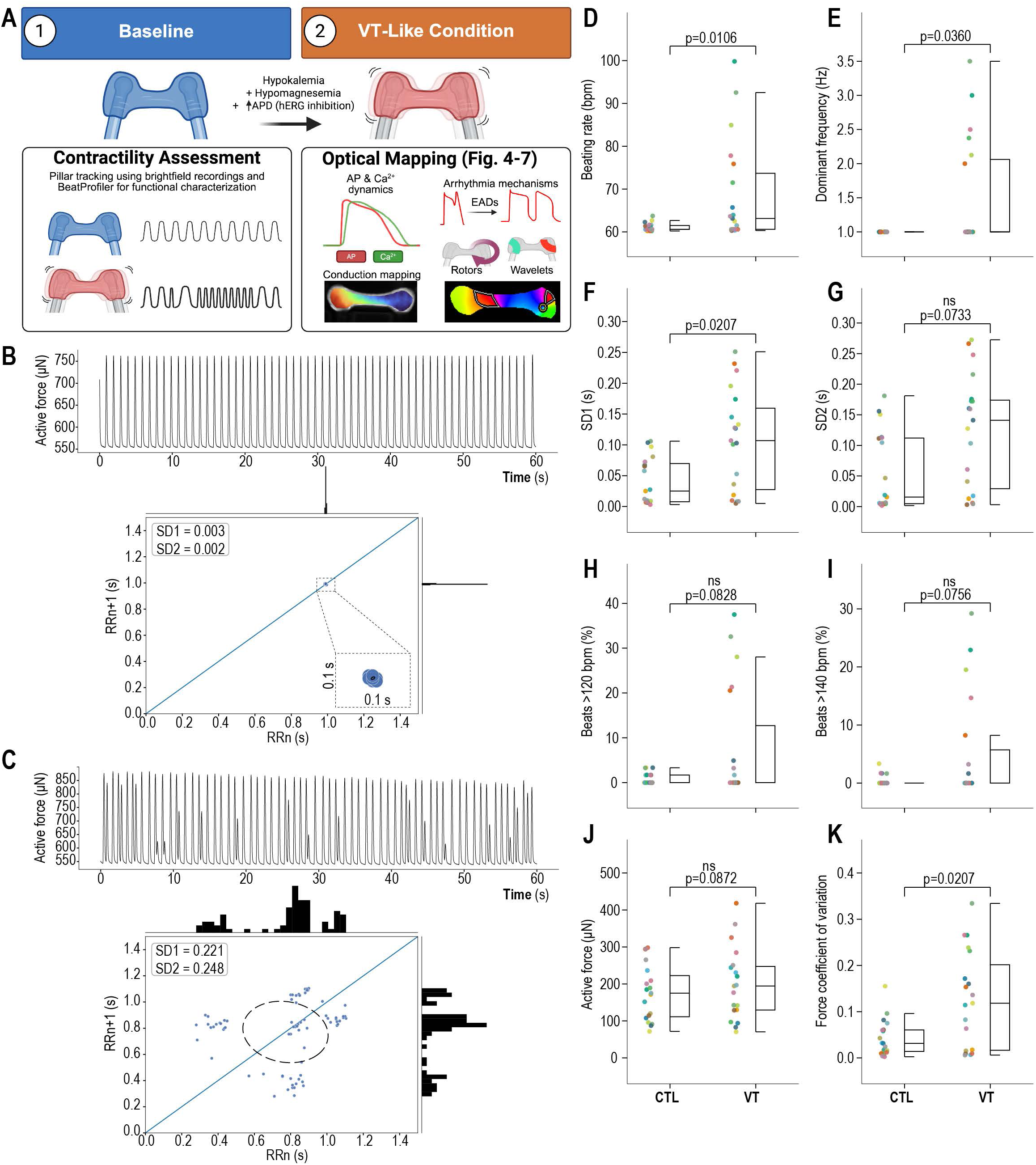
Generation of VT-like condition in EHTs. High-dose E-4031 (500 nM) combined with low extracellular K⁺ and Mg²⁺ created a proarrhythmic substrate that emerged within approximately 1 min and persisted for the duration of the observation period (<5 min). **A**. Diagram of contractility and optical mapping assessments under baseline and VT-like conditions. **B**. Representative active force trace and Poincaré plot from a control (CTL) EHT at 1 Hz pacing demonstrates 1:1 capture and consistent beat-to-beat synchrony. **C**. Representative active force trace and Poincaré plot from an EHT following acute exposure to low K⁺/Mg²⁺ modified Tyrode with 500 nM E-4031 (VT condition) shows transient increases in beating rate and beat-to-beat variability. **D-K**. Summary quantifications of N = 19 EHTs (4 batches) recorded under CTL and then VT conditions. All data are shown as box plots (median, IQR, and 1.5X IQR whiskers) with an overlayed scatter points. Paired data were compared using a two-sided paired Wilcoxon signed-rank test because several metrics showed non-normal paired differences.

Field electrical stimulation delivered by C-Pace (IonOptix) produced regular, synchronous contractions at 1 Hz in culture media (**Fig. 3B**, CTL condition). The same tissues were then exposed to low K⁺/Mg²⁺ plus E-4031 solution, and within 1 min, exhibited periods of rate acceleration beyond pacing frequency, resulting in variable force generation and increased inter-beat variability (**Fig. 3C**, VT condition). We observed significant increases in beating rate, dominant frequency, inter-beat variability (SD1 and SD2 Poincaré analysis), and the proportion of beats >120 and >140 bpm in VT vs CTL condition (**Fig. 3D-I**). While average force generation was preserved in VT, the coefficient of force variation increased significantly likely due to arrhythmic beating behavior (**Fig. 3J-K**). These findings suggest that the combination of hERG inhibition with hypokalemia and hypomagnesemia initiates VT episodes in EHT; therefore, we performed optical mapping studies to dissect the underlying mechanisms.

### 2.6. Optical mapping reveals drivers of VT in EHTs

Dual voltage and Ca²⁺ optical mapping was performed across multiple batches. Due to rapid photobleaching of fluorescent signals and transient nature of arrhythmic phenotypes, initial recordings solely utilized VT-inducing conditions under 1.0 Hz field electrical stimulation to maximize VT episodes recordings. Following protocol optimization, we implemented a baseline condition, recording each EHT while perfusing with normal Tyrode’s solution (CTL) before switching to VT condition, enabling within-tissue comparisons in a subset of experiments (N = 12 EHTs for CTL and N = 13 for VT). To maximize synchronization to pacing within short imaging window (∼19 s) the pacing frequency was increased to 1.5 Hz in some batches. Overall, CTL tissues synchronized to pacing with minor deviations attributable to occasional spontaneous beats (<5%). In contrast, VT episodes induced by low K⁺/Mg²⁺ plus E-4031 frequently accelerated beyond the commanded pacing rate and became irresponsive to pacing (**Fig. 4A**), a response not observed in any CTL tissue. Pooled data across pacing frequencies show increased percentage of time beating at >120/140 beats per min in the VT vs CTL condition (**Fig. 4B**). As arrhythmic behavior under VT condition was largely frequency-insensitive, subsequent quantifications are hereinafter presented as CTL and VT.

**Figure 4.**
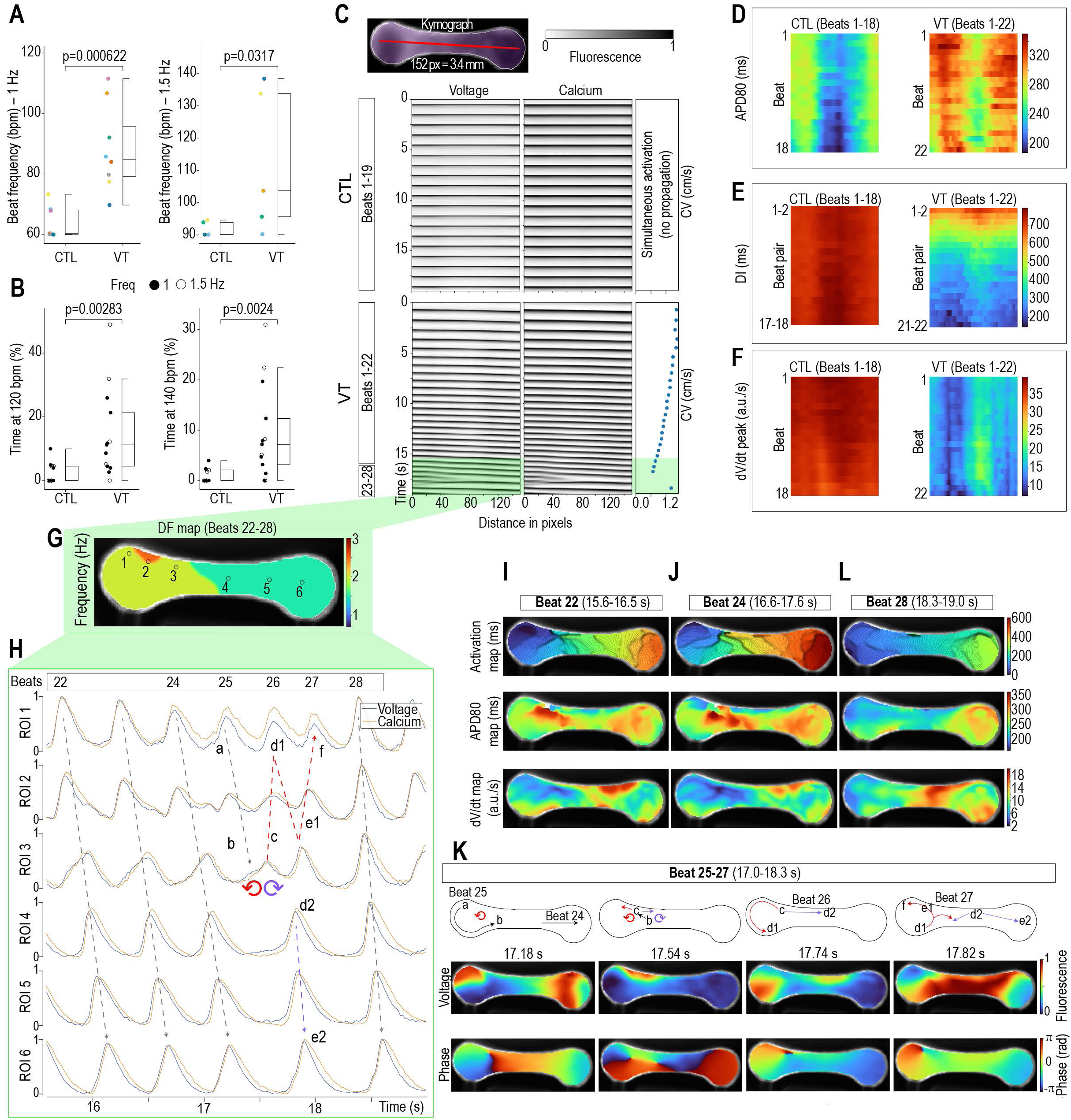
VT condition prolongs APD_80_, increases dispersion and promotes wavebreak and reentry in EHTs. Multiple long (19 s) dual voltage and Ca²⁺ optical mapping recordings were obtained per condition at 1.0 or 1.5 Hz field electrical stimulation, recreating tachyarrhythmia observed in contractility assays. **A**. Beating frequency at 1.0 and 1.5 Hz field electrical stimulation. **B**. Fraction of recording time spent above 120 and 140 bpm for EHTs paced at 1.0 Hz (●) and 1.5 Hz (○). Shapiro-Wilk indicated non-normality for several distributions; unpaired two-sided Mann-Whitney U test were applied (CTL N = 12, VT = 13, 4 batches). **C**. Voltage (left) and Ca²⁺ (right) kymographs from a CTL (top) and VT (bottom) recording showing simultaneous activation under pacing in the CTL and progressive loss of capture, beat acceleration, and slowed CV under VT (**Suppl. Video 5-6**). **D-F**. Heatmaps showing pixel-wise changes across beats for APD_80_, DI, and dV/dt from a representative EHT under CTL (left) and VT (right) conditions. APD_80_ becomes prolonged and spatiotemporal localized heterogeneity increases under VT condition, while DI shortens and dV/dt slows, likewise becoming more heterogenous. **G**. Dominant frequency (DF) maps from beats 22-28 outlined in green highlights regions of increased frequency leading to wavebreak and reentry across the EHT. **H**. ROI-based voltage and Ca²⁺ traces transition from slow but continuous propagation to slowed conduction causing conduction block and reentry before returning to slow but continuous propagation. Activation, APD_80_, and dV/dt maps for beats 22 (**I**) and 24 (**J**) show decreased CV and increased spatiotemporal APD_80_ heterogeneity, producing dV/dt minima and conduction delays preceding wavebreak. **K**. Diagrams (top), voltage fluorescence (middle), and phase (bottom) maps for beats 25-27. Signal emerge from the left head, move in a downward circular motion (a-b), slowing down within the head-neck space (b-c), bifurcating onto areas that exhibited low APD (prior beat, **J**), while avoiding high-APD areas. Two PS/rotor arise: one moving leftward initiating a second rotation in the left head (c-d1) and the other propagating on the upper side towards the right head (c-d2). After rotation, the left wave (d1-e1) collided with a bidirectional wave initiated near the right neck (from d2), and both wavefronts extinguished short after. These events lead to the restoration of continuous but slow propagation by beat 28 (**L**), the dissipation of APD gradients, and the normalization of dV/dt (**K**).

Kymographs were used to evaluate beat-resolved electrophysiology and propagation dynamics across the longitudinal axis in CTL and VT recordings. Representative CTL recording show activation in straight lines, consistent with synchronized capture throughout the entire longitudinal axis due to field stimulation (**Fig. 4C** and **Suppl. Video 5**). After switching to low K⁺/Mg^+2^ plus E-4031, 9/13 tissues progressively lost pacing capture, exhibited acceleration and decelerating burst, and developed spatiotemporally fragmented activation consistent with wavebreak and transient reentry (**Fig. 4C**). Spatial APD heterogeneity can promote wavebreak and occurs when the AP wavefront fails to propagate in regions of cardiac tissue that have not fully recovered from a previous excitation. When wavebreak occurs in a localized area, it can lead to reentry and fibrillation.^25^ The representative VT recording shown in **Fig. 4C** displays an activation starting from the top left corner of the EHT that progressively slows down (CV from 1.6 to 0.7 cm/s) as beating frequency accelerates before wavebreak and partial recovery towards the end of the recording (**Suppl. Video 6** shows events occurring between 15-19 sec). Kymographs were used to quantify beat-to-beat changes in APD_80_, diastolic interval (DI), and action potential upstroke departure velocity (dV/dt), which are presented as heatmaps with distance along the tissue on the x-axis and successive beats on the y-axis. Under CTL conditions, EHTs showed a clear APD gradient that is prolonged towards the heads, but stable beat-to-beat APD_80_, consistent with 1:1 capture during field electrical stimulation (**Fig. 4D**, left panel). The VT condition, in contrast, displays APD prolongation with more severe regional APD_80_ gradients that evolved over the recording period (**Fig. 4D**, right panel). DI and dV/dt, which are highly consistent under CTL conditions, become spatially and temporally variable in the VT condition (**Fig. 4E-F**). Across recordings, conduction block consistently occurred in regions exhibiting pronounced APD prolongation with depressed dV/dt, positing repolarization delay as a potential mechanistic substrate for the functional conduction barriers observed under VT conditions.

Dominant frequency mapping during activation fragmentation (beats 22-28 in the representative VT recording) revealed regions of localized high-frequency activity towards the left EHT head, where frequencies rose to 2-2.5 Hz (**Fig. 4G**). Voltage and Ca²⁺ traces in ROIs across the frequency gradient demonstrate a transition from organized wavefront propagation to regions of conduction block, accompanied by transient phase singularities (PS) and rotors during wavebreak-associated beats (**Fig. 4H** and **Suppl. Video 6**). Activation maps show progressive delays in activation times (from ∼200 during initial beats to ∼600 ms on beat 24) with larger APD_80_ gradients around the left EHT head. These changes coincided with local minima in dV/dt around the left EHT head on the beat preceding conduction block, wavebreak, and rotor formation (**Fig. 4I-J**). During wavebreak, AP propagation avoided areas that had longer APD_80_ and low dV/dt (**Fig. 4J**) resulting on a rotor in the left head, whereas on the right, propagation entered the EHT shaft, generating a beat that merged with the left-head rotor (**Fig. 4K**). By beat 28, a more organized propagation pattern was restored with faster activation, and dissipation of APD_80_ and dV/dt gradients around the left EHT head (**Fig. 4L, Suppl. Video 6**).

### 2.7. Phase singularity tracking confirms transient rotor formation exclusively in VT EHTs

EHTs in the VT group exhibited recurrent, transient rotors that promoted reentry and further increased beating frequency. In some recordings, these rotors were scarce and short-lived (as described in **Fig. 4K**), whereas in other recordings multiple rotors were observed leading to local increases in beating rate (**Fig. 5**). Kymographs from another representative VT EHT (**Fig. 5A**) show a fragmented beating pattern along the longitudinal axis, that coincides with PS/rotors on both the left and right EHT heads. Initially, APD_80_ was substantially longer but homogeneous in the right head, with persistent gradients across the tissue (**Fig. 5B**). Interestingly, the right EHT head exhibited local APD gradients like the ones observed in **Fig. 4** prior rotor initiation, whereas the left head exhibited APD alternans (alternating short-long APD) together with local APD gradients prior rotor initiation. DI shortened substantially during beat-to-beat spontaneous acceleration from ∼764 to ∼201 ms on the right side and ∼261 ms on the left side immediately before reentry (**Fig. 5C**). Progressive regional reductions in upstroke velocity dV/dt observed first around right EHT neck and then around left EHT neck preceded rotor formation (**Fig. 5D**).

**Figure 5.**
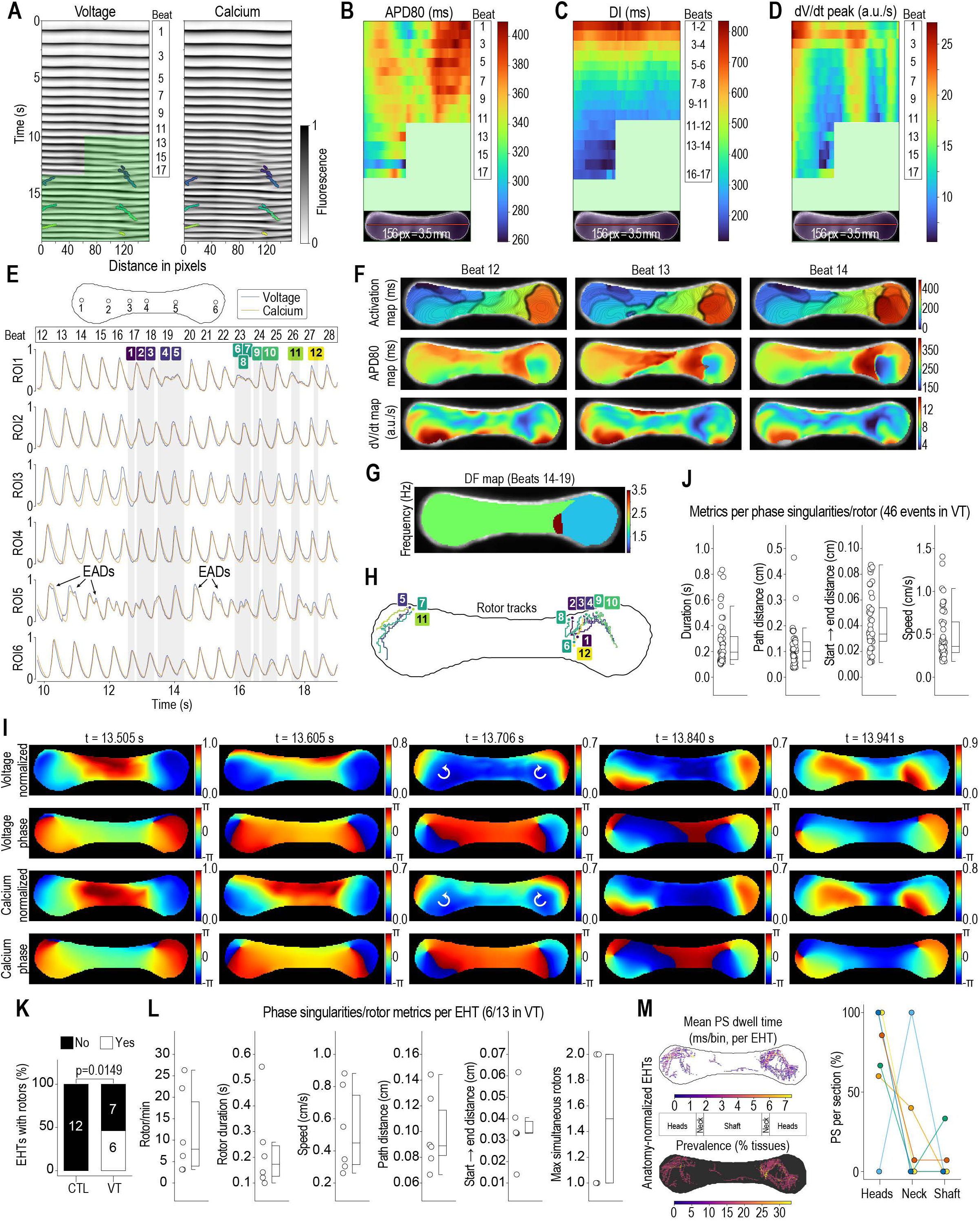
Phase singularities and rotor dynamics in VT-condition EHTs. **A**. Voltage (left) and Ca²⁺ (right) kymographs with PS tracks overlay from a second representative VT recording. Continuous propagation was analyzed using kymographs (**B-C**), and areas with signal fragmentation were masked (green) and resolved with ROI-based trace plots (see below). **B**. Pixel-wise longitudinal APD_80_ heatmap demonstrates APD gradients between EHT heads and shaft as described in Fig. 4C. Prior to reentry APD dispersion within the heads progressively increased as DI progressively shortened during spontaneous beating acceleration (**C**). Areas with reduced excitability (dV/dt minima) increased over time near the EHT head-neck joints and preceded the formation of PS/rotors. **E**. ROI-based voltage (blue) and Ca²⁺ (orange) traces obtained from 6 ROIs between 10-19 s (masked green area) show APD prolongation with EADs in the areas preceding rotor formation. Gray shading represents the times when PS were detected using a PS-tracking algorithm. **F**. Activation time, APD_80_, and dV/dt maps for beats 12-14 show a progressive increase in activation time and circular propagation around the right EHT head. APD_80_ increased over time and progressively generated high-low APD areas around the right EHT head, linking the initial slow and circular propagation to a reduction in dV/dt that preceded conduction block, wavebreak and reentry. **G**. Dominant frequency maps show areas beating at 3.5 Hz in the right EHT neck-head joint caused by reentry driven by rotor formation in the right EHT head. **H**. PS trajectories observed during the recording in the EHT and shown in **A** and **E** overlays. **I**. Sequential voltage (top) and Ca²⁺ (bottom) frames with phase maps show simultaneous rotors in the left (counterclockwise) and right (clockwise) EHT heads, generating rotating wavefronts that converge at the center (**Suppl. Video 7**). **J**. Summary quantifications of all PS events (46 total) across N = 13 VT EHTs from 4 batches. **K**. Proportion of CTL tissues with rotors (0/12) versus VT (6/13). Fisher’s exact test was used. **L**. Summary quantifications of PS/rotor events in the 6 rotor-positive VT EHTs. **M**. Anatomy-normalized overlay of PS dwell time and regional prevalence shows predominant localization in the tissue head and neck regions. All quantifications except panel **M** were reported with box plots (median, IQR, and 1.5X IQR whiskers) with overlaid scatter points. In panel **M**, points from the same EHT are connected across head, neck, shaft regions.

ROI-based traces and beat-resolved maps show a progressive slowing of CV and a relocalization of the propagation signal in a concentric pattern around the right EHT head that precede chaotic wave propagation at ROI 5 (**Fig. 5E-F, Suppl. Video 7**). Interestingly, considerable APD_80_ dispersion in this area progressively increased (**Fig. 5F**) leading up to rotor formation, with longer APD regions exhibiting dV/dt minima around the right EHT neck. APD prolongation was likely caused by early afterdepolarizations (EADs) that preceded rotor formation around the right head region (**Fig. 5E**); a similar finding was observed in **Fig. 4H** (beats 22-23, ROI2) with EADs preceding rotor formation.

Dominant frequency maps of beats 14-19 show localized increases in beating frequency, reaching 3.5 Hz in a narrow area of the right EHT neck, due to reentry driven by rotors around the right head that returned to this point (**Fig. 5G**). Multiple rotor trajectories were calculated by detecting PS with Hilbert-transform calculations and are shown in **Fig. 5H** and overlapped on ROI-based traces (**Fig. 5E**). Half rotations started earlier on beat 14 and consolidated on PS on beat 17 (∼12.5 s) with some PS coexisting on both heads of the EHT (**Fig. 5I**)

Across all VT EHTs, 46 PS/rotor events were detected, with a median duration of ∼200 ms, path length of ∼100 μm, start➔end distance of ∼30 μm, and speed of ∼0.3 cm/s (**Fig. 5J**), indicating localized, slow, and short-lived rotor formation. Spatiotemporally conserved rotor termination and initiation events near the tissue borders may indicate the presence of longer-lasting rotors with trajectories beyond mapped EHT regions and are addressed in the discussion. Summary tissue statistics show that rotors were exclusively observed under VT conditions, with 6 of 13 EHTs under VT conditions exhibiting rotors, whereas no rotors were observed under CTL conditions (**Fig. 5K**). As expected, most of the detected rotors were short-lived (∼0.2s) and covered short distances within the EHTs (∼0.1cm; **Fig. 5L**). When tissue geometry is normalized using principal component alignment, PS dwell-time heatmaps reveal enrichment of PS within the head and neck segments with travel paths frequently following head curvature (**Fig. 5M**).

### 2.8. Triggered activity accompanies VT initiation and dynamically interacts with reentry

EADs commonly co-localized with regions of APD dispersion and dV/dt depression immediately before rotor formation (as shown in **Fig. 4-5**), suggesting that triggered depolarizations may initiate tachyarrhythmias and contribute to the functional conduction barriers that stabilize reentrant activity. In addition to reentry, 4/13 VT tissues exhibited single beat triggered activity driven by EADs, most frequently during late repolarization. Only one of these tissues exhibited rotors and progressive acceleration. No EADs were observed in CTL tissues. Voltage and Ca²⁺ traces from a representative EAD-positive EHT are shown in **Fig. 6A**, with propagation direction and activation maps for the beats immediately before and with EADs highlighted in **Fig. 6B**. EAD-dependent triggered activity propagated from head-to-head or locally, colliding with wavefronts coming from the other head.

**Figure 6.**
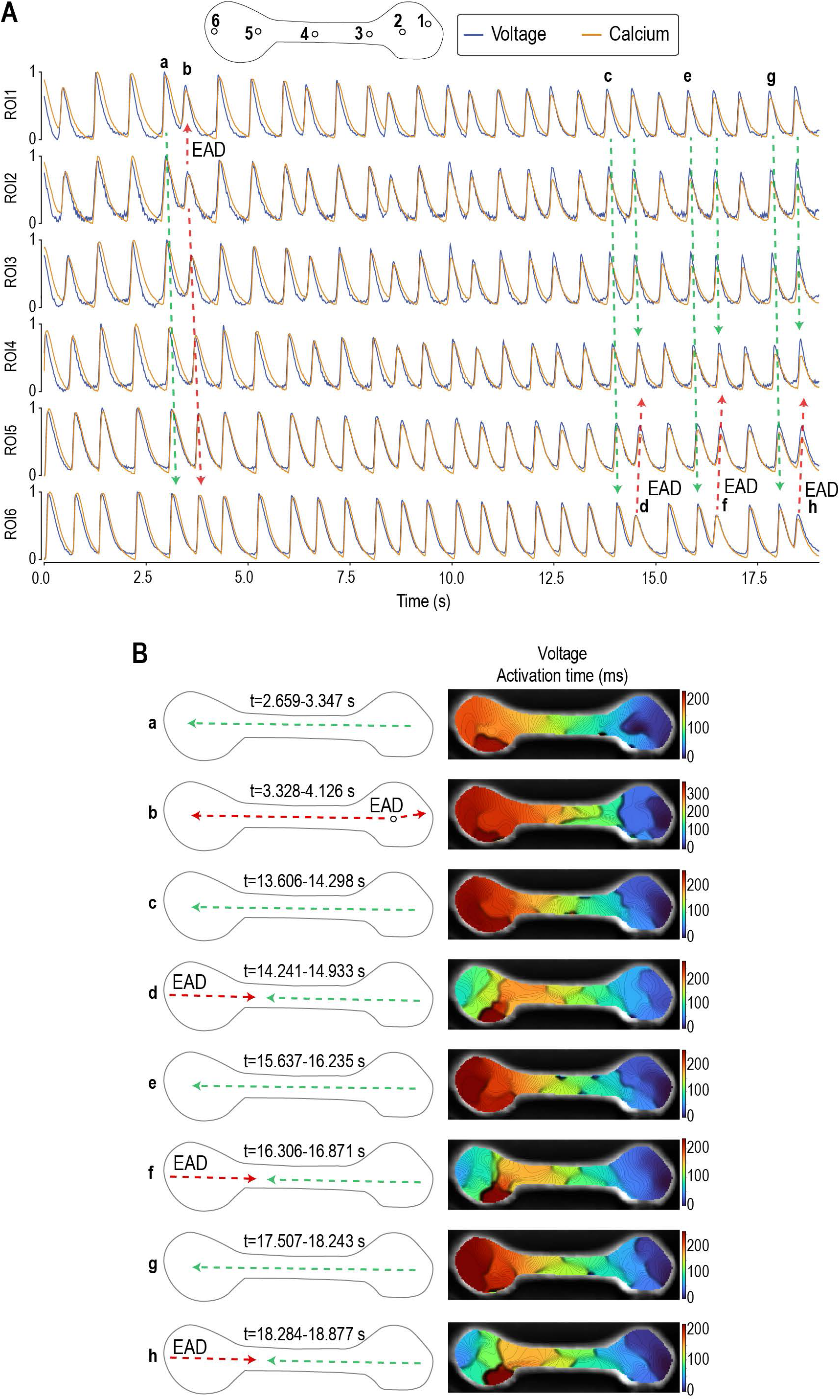
EADs contribute to regions of functional conduction block in VT-condition EHTs. **A**. Voltage (blue) and Ca²⁺ (orange) traces from a representative VT-condition EHT highlight phase-3 EADs (red dotted arrows) and the “normal” propagation immediately before and coinciding with the EAD (where applicable, green dotted arrows). EAD-driven triggered activity was observed in 4/13 tissues and was associated with increased beating rate or multi-foci activation. Phase-2 and phase-3 EADs were detected in 1/13 and 3/13 EHTs, respectively, yielding 26 and 2 total triggered-activity events across recordings. **B**. Schematic representation of propagation direction for the marked beats from panel A (left), with corresponding voltage activation maps (right) showing EAD-driven triggered activity and bifocal activation at beats d, f, and h.

### 2.9. Does VT progresses to VF in EHTs?

While rare (2/13 of VT EHTs), transient episodes of short-lived highly complex activation patterns characterized by the presence of multiple wavelets and PS were observed, with partial AP and Ca²⁺ wavefront segmentation consistent with chaotic signal resembling VF-like episodes (**Fig. 7**). In one of these tissues, voltage and Ca²⁺ kymographs with overlayed PS traces demonstrate spatially separated simultaneous activation at opposite ends of the tissue generating up to 3 wavefronts or wavelets (**Fig. 7A-B, Suppl. Video 8**). These wavelets and PS coexisted simultaneously in this tissue over the entire recording (**Fig. 7C-D**) but resolved quickly in a more organized VT phenotype. Another example is shown in **Fig. 5G-I**, where two PS generated full rotations on both EHT heads.

**Figure 7.**
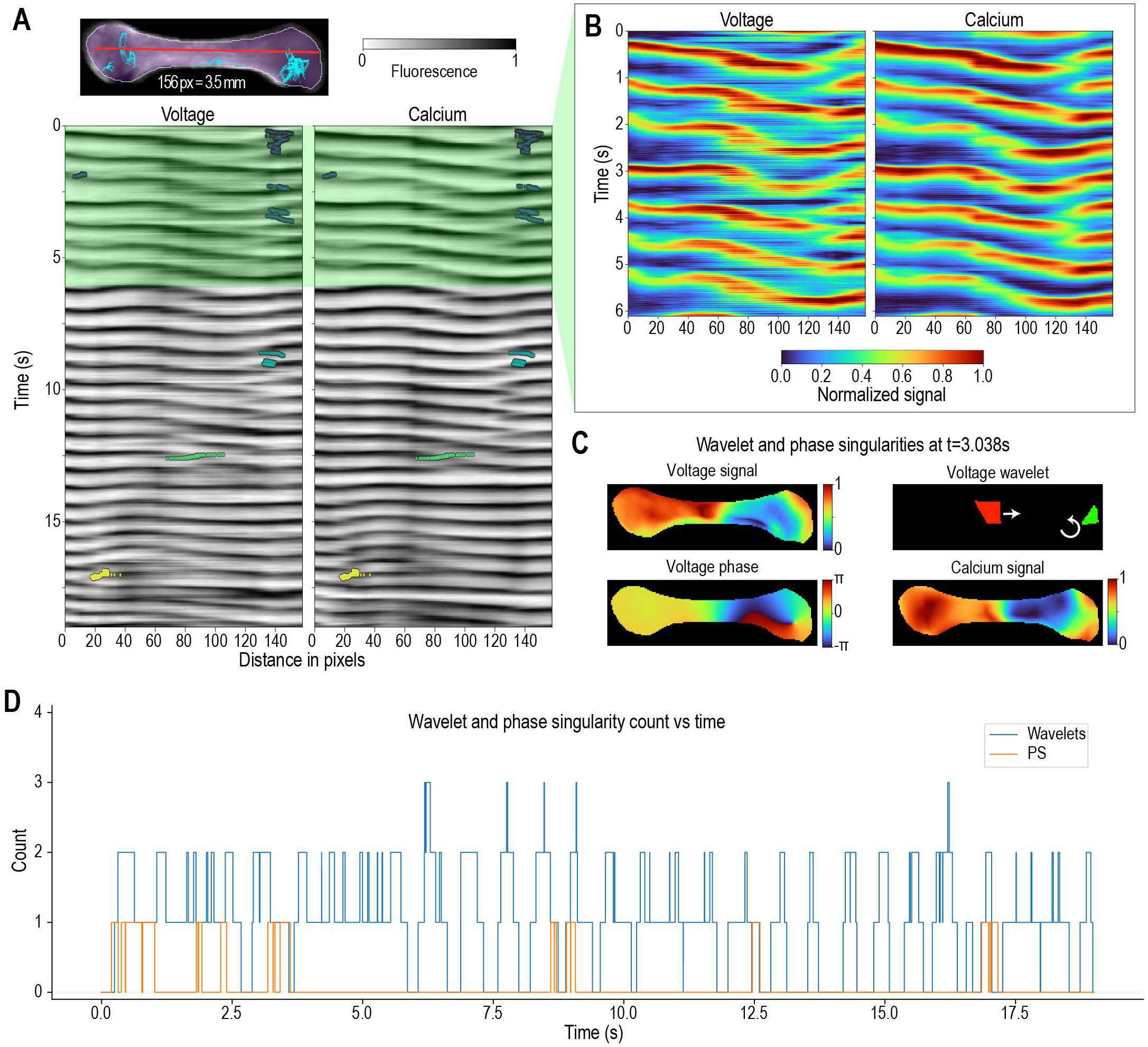
Highly complex activation patterns in a subset of VT-condition EHTs exhibit VF-like fragmentation. **A**. Voltage (left) and Ca²⁺ (right) kymograph with overlaid PS tracks from a VT EHT. **B**. Magnified view highlights the spatiotemporal heterogeneity of voltage and Ca²⁺ signals (**Suppl. Video 8**). **C**. Voltage signal and phase map with highlighted voltage wavelets reveal abnormal activation coinciding with a PS and two simultaneous wavelets and signal heterogeneity. **D**. Quantification of simultaneous wavelet and PS counts over time in the representative VT EHT demonstrates VF-like presence of multiple wavelets.

## 3. Discussion

Non-animal, human iPSC-derived EHTs are gaining traction as translational platforms, yet their capacity to reproduce adult ventricular arrhythmia mechanisms remains unclear due to size limitations and tissue immaturity. To date, no study has provided high-resolution tissue-level spatiotemporal electrophysiology to determine whether these models recapitulate adult electrophysiology and test whether classic initiators and drivers of VT/VF manifest in them. We therefore adapted an open-source EHT platform and implemented optical mapping that achieved temporal and spatial resolution previously attainable only in whole hearts. Baseline measurements showed restitution curves and AP-CaT coupling comparable to adult hearts. By imposing a pro-arrhythmic condition, we reproduced the sequence of substrate evolution underlying VT/VF: localized APD dispersion, loss of excitability, conduction block, wavebreak, and rotor-mediated reentry. Mapping revealed that repolarization instability creates transient “physical barriers” that anchor rotors, and we identified anatomical hotspots serving as arrhythmic foci. Our work demonstrates that mature EHTs can recapitulate adult electrophysiology and provide a robust non-animal platform for mechanistic arrhythmia studies. The ability to quantify electrophysiological alterations and detect major electrical events described here can be applied to other diseases that increase arrhythmogenic susceptibility.

### 3.1. Electrophysiological maturation of EHTs

The measured baseline APD_80_ (∼430 ms at 1000 ms CL to ∼269 ms at 333 ms CL) fall within the normal adult human range (∼317-360 ms at 1000 ms CL^26^; ∼300/400 ms at 500/1000 ms CL^27^; ∼220/400 ms at 333-2000 ms CL^28^). Longitudinal CV averaged ∼39 cm/s and fell to ∼21 cm/s at fastest pacing, values that sit within the lower end of human adult ventricular conduction (56 cm/s longitudinal, 14.6 cm/s transverse in donor heart wedges^29^; 95 cm/s longitudinal and 45 cm/s transverse in non-failing human hearts^28^; and 45-67 cm/s in large animal models^30^). AP-CaT latency matched adult mouse and rabbit heart values^31–33^, suggesting mature and efficient ECC. Pharmacological validation with the hERG blocker E4031 (100 nM) prolonged APD_80_, confirming faithful recapitulation of species-specific ion channel modulation. Collectively, these data demonstrate that millimeter-scale EHTs can reproduce adult-like tissue-level electrophysiology by 1 month.

### 3.2. Pro-arrhythmic substrate generation and mechanisms

Acute high-dose hERG blockade with 500 nM E-4031 under low K⁺/Mg²⁺ created a proarrhythmic substrate that emerged within approximately 1 min and persisted throughout the observation period (<5 minutes). High-dose E-4031 alone further prolonged APD but did not induce VT-like arrhythmias, consistent with rabbit heart data where APD prolongation alone does not create a pro-arrhythmic substrate. When combined with low K⁺/Mg²⁺, E4031 produces heterogeneous repolarization that leads to VT/TdP.^19,33^ In this model, arrhythmias emerge from a combined failure of repolarization reserve and triggered substrates, where low extracellular K⁺ potentiates hERG blockage^34^, reduces I_K1_ currents, and inhibits Na⁺/K⁺ ATPase leading to repolarization heterogeneity and spontaneous ventricular firing ^35^, and low extracellular Mg²⁺ increases L-type Ca²⁺ channel (LTCC) currents and SR Ca²⁺ release, amplifying afterdepolarization propensity in ventricular myocytes.^36^

The prevailing mechanistic view underlying VT/TdP in low K⁺/Mg²⁺ plus 500 nM E-4031 has historically centered on EADs. Previous observations suggest that spontaneous Ca²⁺ events can precede voltage changes at EAD initiation sites, supporting a Ca²⁺-overload contribution to EAD generation and polymorphic activity.^18^ Building on this, Maruyama et al. reported that this phenomenon is dominated by phase-3 EAD-mediated triggered activity arising at borders between long- and short-APD regions; in some TdP episodes, they observed EAD-induced initiation of a reentrant rotor with a drifting core, emphasizing that APD gradients can couple triggered activity to reentry.^37^ More recently, Alexander et al. proposed that triggered activity leading to premature ventricular contractions and TdP is better explained by LTCC reactivation originating in discrete regions with dynamically generated, extreme long-short APD gradients, rather than by EADs.^18^ They further suggested that low levels of LTCC blockade can suppress this activity.

Our observations are consistent with these mechanisms. We observed phase-2 and phase-3 EADs, resulting in 26 and 2 total triggered activity events in 1/13 and 3/13 of the studied EHTs. This agrees with previous data showing that 74% of triggered activity is driven by phase-3 EADs in rabbit hearts.^19^ However, in most recordings (9/13), EHTs exhibit progressive acceleration and deceleration, starting from single or multiple foci, with extremely slow propagation. Prepotential slopes preceding the AP upstroke were observed in multiple recordings, which is in agreement with an increase in automaticity likely caused by I_K1_ blockage and partial membrane repolarization due to impaired Na⁺/K⁺ ATPase under hypokalemic conditions. No spontaneous increases of Ca²⁺ were observed during AP prepotentials, discarding the possibility of DADs. As tissue started to accelerate, optical mapping showed an increased localized dispersion of APD_80_, accompanied by a progressive shortening of DI and a marked reduction in upstroke velocity, producing spatially heterogeneous repolarization gradients. These changes slowed conduction locally, promoted wavebreak, and enabled the emergence of short-lived PS that manifested as focal rotors. Rotor trajectories were confined largely to the head-neck junctions, regions where APD was longest and dV/dt lowest, suggesting that a combination of prolonged repolarization and impaired excitability is sufficient to generate re-entrant circuits. The observations are consistent with classical mechanisms of ventricular tachycardia in which steep APD gradients^38^ promote conduction block, wavebreak and reentry. Intriguingly, sudden EAD formation preceded conduction block in areas with high and low APD gradients, leading to “physical barriers” for wavebreak and rotor formation. Recent tissue-scale modeling suggests that EADs in isolated cells are highly variable due to stochastic processes, leading to beat-to-beat APD variability, however in cardiac tissue the fluctuations are largely dampened. EADs manifest as discontinuous alternans that lead to pronounced spatial heterogeneities which are strongly arrhythmogenic, as the sudden onset of EADs on alternate beats in cardiac tissue promotes conduction block and reentry.^37^

### 3.3. PS/rotor localization in EHTs

The concentration of PS/rotors near the tissue heads/neck implicates EHT morphology and regional heterogeneity in the development of arrhythmogenic susceptibility that could arise from mechanical strain gradients, fiber orientation, tissue thickness, among other variables. In adult hearts, PS clustering occurs in a nonrandom spatial distribution and colocalizes with normal anatomic heterogeneities along epicardial vessels, ridges of endocardial trabeculae, and papillary muscle insertions.^39^ These locations corresponded to areas of fiber apposition with different angulations and to intramural vessels. Similarly, rotational anisotropy and tissue boundaries play an important role in generating breakthrough waves during VF.^40^ Mechanistically, fiber rotation induces curvature in scroll-wave (3D rotor) fronts, which weakens conduction and further facilitates wave break.^41^ Additionally, dispersion of mechanical strain and electrical recovery have been described, underscoring mechanoelectrical feedback as an important part of arrhythmogenesis^42^, which can induce deterioration of stable spiral waves into turbulent wave patterns.^43^ Interestingly, we observed APD gradients between heads and shaft of EHTs which under normal conditions did not trigger arrhythmic events and only generated rotors when a pro-arrhythmic substrate was in place: APD dispersion within the head/neck leading to long- and short-APD regions within the EHTs neck. These observations are consistent with anatomically realistic 3D ventricular models that incorporate physiological fiber orientation, which show that dynamical instability of propagation (e.g., restitution-driven instability) is the primary trigger of wavebreak and VF. Cardiac geometry and anisotropic fibers act as modifiers that lower the breakup threshold and shape filament folding, drift, and surface interactions, thereby amplifying the likelihood and complexity of VF once instability is present. Conversely, when dynamical instability is sufficiently suppressed (e.g., by flattening restitution), organized electrical activity can persist despite structural heterogeneity. Thus, anatomy shapes fibrillatory dynamics but does not alone generate sustained VF without an unstable electrophysiological initiator.^44^

The short-lived VF-like episodes observed in 2/13 VT/EHTs are notable as they display a surprisingly complex pattern (2-3 coexisting wavelets with intermittent phase singularities; **Fig. 5,7**), indicating that proarrhythmic stressors can transiently drive multi-wavelet dynamics even in EHTs. However, the short duration argues against sustained VF and supports wavebreaks that are generated but not maintained. Our EHTs are constrained to ∼4 mm in length, which may be insufficient to reproduce fibrillatory patterns due to space constraints in accordance with the critical mass hypothesis^45^, and it is possible that anatomical barriers along the transverse axis (∼0.5 mm) or at head-shaft transitions promote rotor annihilation around EHT heads. Moreover, for a rotor to remain stable, a critical minimum inter-PS distance of approximately 4 mm is required during fibrillation in sheep hearts; if singularities of opposite chirality move closer than this threshold, they mutually annihilate.^46^

### 3.4. Limitations and future directions

Several limitations of the present work should be considered. Although our data suggest that millimeter-scale EHTs begin to reproduce adult-like, tissue-level electrophysiology by ∼1 month, our study was not designed to define the full maturation trajectory. We did not perform longitudinal studies of maturation including imaging, ultrastructure and other techniques widely used to compare human myocardium and engineered constructs. Instead, we focused on baseline electrophysiological characterization at 1 month and VT experiments at 2 months. Our findings provide a foundation for more detailed, time-resolved studies of electrophysiological maturation with longitudinal designs spanning multiple timepoints.

We used low K⁺/Mg²⁺ plus E4031, as a model of acquired LQTS. Our findings may not be applicable to other types of LQTS or VT. Optical mapping provides a top-surface weighted readout, and events that exit or occur beyond the field of view cannot be captured. EADs driving triggered activity usually emerged from areas around the heads (**Fig. 6**) and it is possible that they could emerge from below the imaging field. PS and rotor criteria were accordingly and intentionally conservative to reduce duplicate event identification which may have underestimated PS and rotor event durations (**Fig. 5**). Although some tolerance was applied to merge events (within 5 pixel and a maximum 10 ms gap), longer padding created artificial trajectories connecting rotors from opposite heads. Fluorescent dyes and electromechanical uncoupling are known to influence cardiac electrophysiology and could also add confounding factors. Though we attempted to minimize these effects by optimizing dye loading protocols, this limitation underscores the value of orthogonal validation strategies where feasible. Finally, optical mapping systems are expensive which may limit the applicability of the electrophysiological characterization described here. Open-source alternatives, however, do exist and the CMOS-based approaches can partially mitigate this barrier.^47–49^ Future investigations may improve upon our work by leveraging the apparent impact of EHT anisotropy and morphology, as well as evaluating interventions to normalize repolarization or excitability under pathophysiological conditions to better understand the mechanisms of human cardiac arrhythmogenesis.

### 3.5. Concluding remarks

In summary, our findings establish that millimeter-scale, human iPSC-derived EHT can faithfully reproduce the electrophysiological hallmarks of adult ventricles (restitution curves, conduction velocities, AP-Ca²⁺ coupling). By imposing a clinically relevant low K⁺/Mg²⁺ along with hERG blockade we induced the full cascade of pro-arrhythmic substrate evolution: localized APD dispersion, excitability loss, conduction block, wavebreak, and the emergence of rotor-mediated reentry. The spatial confinement of PS underscores how geometric heterogeneities can serve as physical anchors for rotors when repolarization reserve is compromised. Collectively, these data demonstrate that EHTs constitute a robust, non-animal system for dissecting the mechanistic underpinnings of ventricular tachyarrhythmias and for pre-clinical screening of anti-arrhythmic strategies. Although the finite size of our construct limits sustained VF, we evaluated multiple parameters, such as APD dispersion and areas of long-short APD, that precede rotor formation and can be assessed for drug discovery. Future work will expand tissue dimensions, integrate physiologic fiber orientation and mechano-electrical feedback, and couple this optical mapping framework with spatial biology and genome-edited disease models for precision arrhythmia research modeling.

## 4. Methods

### 4.1. Cell sources and culture

Human iPSC-ventricular cardiomyocytes (hiPSC-CMs) were derived from the hPSCreg line #TMOi001-A (Gibco Episomal iPSC Line; A18945), differentiated and expanded according to the massive cardiomyocyte expansion protocol.^50^ Only batches that formed confluent, spontaneously beating monolayers after four days of metabolic selection were used for further expansion. Cells underwent two passages in the presence of 2 μM CHIR-99021 (Selleckchem S2924) for up to 35 days, followed by a one-week recovery period without CHIR-99021. Our cardiomyocyte population was >95% pure. Primary human ventricular fibroblasts (Lonza NHCF-V) were expanded per manufacturer protocol and used up to passage 5.

### 4.2. Silicone-form generation

milliPillar designs^17^ were 3D-printed with temperature-resistant resin (Phrozen Sonic Mini 8K, TR300 Ultra-High Temp 3D Printing Resin). Several modifications were introduced to the computer numerical control (CNC) milliPillar molds ^17^ to improve printing accuracy: (i) all modules were interconnected to stabilize the individual milliPillar units; (ii) the mold faces surrounding each milliPillar were raised by 200 μm, preserving pillar geometry while allowing flat sanding of both faces with 2000-grit sandpaper; (iii) registration pins were replaced with M2X6mm screws, (iv) the top pocket microchannel was eliminated from the top mold and replaced by a fence to facilitate handling of individual milliPillar modules. To prevent cure inhibition, printed molds were treated with 405-nm UV light for 1h while rotating continuously using a 3D printing curing station, followed by heat treatment at 100 °C for 3 h and subsequently at 65 °C overnight. After two casting cycles, cure inhibition was no longer observed.

Ease Release 200 (Mann Release Technologies) was sparsely sprayed on mold faces to facilitate mold release. Polydimethylsiloxane (PDMS) (base-to-curing-agent 10:1, EMS 24236-10) was degassed, poured into the printed molds, degassed again, then clamped on both faces and cured for 24 h at 65 °C as described.^17^ After unmolding, molds were cured for additional 24 h, washed with MilliQ water, 70% ethanol, and MilliQ water, then dried at 65 °C. The milliPillar modules were trimmed, tapped onto a very thin PDMS layer and glued into 6-well plates, then cured overnight at 45 °C.

Each glued milliPillar module was plasma-treated using a handheld corona plasma treater (Electro-Technic Products BD-20AC) equipped with a spring tip electrode to deliver corona plasma into the glued milliPillar module. A total of 15 seconds with the dial set at low-to-medium intensity was enough to increase PDMS hydrophilicity. The plate was subsequently sterilized with 70% ethanol for 30 minutes, air dried, and UV-C irradiated (UV Clave, Benchmark Scientific) for 20 minutes directly under UV bulbs (600 mJ/cm^2^) inside a biosafety cabinet. UV-C indicator cards were used to monitor UV-C exposure (B1450-IC, Benchmark Scientific). After sterilization, milliPillar modules were blocked with KnockOut™ Serum Replacement (KOSR, Gibco 10828-028) for 1h at 37 °C.

### 4.3. EHT plating

Cells were detached with TrypLE Select (Gibco 12563011), centrifuged (300 xg, 3 min), counted, and resuspended in plating medium RPMI (Gibco 11875093) supplemented with B27 (Gibco 17504044), 10% KOSR, 1X penicillin/streptomycin (Gibco 15140122), and 5 µM Y-27632 (Selleckchem S1049). We plated EHTs in batches of at least 12, including a minimum of 3 additional tissues to account for pipetting losses. Each EHT contained 550,000 viable cells at a 3:1 cardiomyocyte to fibroblast ratio. Cells were pelleted and resuspended in 15 µL of matrix solution per EHT, consisting of 12 µL of PureCol® EZ Gel (BICO 5074, 5 mg/mL bovine atelocollagen) and 3 µL plain RPMI supplemented with 25 µM Y-27632, to achieve a final collagen concentration of 4 mg/mL and 5 µM Y-27632.

EHT suspension (15 µL) was homogeneously dispersed without introducing bubbles to each milliPillar module pocket near the pillar heads and by sweeping the pipette tip across the pocket while dispensing to the opposite pillar. We spread the mixture around the upper fence to make sure the mixture remains around pillar heads by capillarity. Here we identified several problems that affected our plating: (i) excess PDMS used to glue inserts was detached by pipet motion during plating, incorporating PDMS into the EHT matrix; (ii) leakage between PDMS and 6-well plate bottom. These problems were solved, as described above, by spreading a very thin layer of PDMS on a flat surface of a 6-well plate and then tapping the milliPillar onto the 6-well plates. By looking under the base of the plate, we were able to determine a continuous layer of PDMS was around milliPillar module, without leaking into the pocket chamber; (iii) bubble incorporation, causing EHTs structural defects. Here, it is important to gently pipet the suspension without introducing bubbles. Plasma treatment significantly reduced bubble generation from air entrapment.

After plating, ∼25 µL plating medium was added along the well wall to prevent dehydration before incubation for 1 h at 37 °C. Plating media was added after, on the sides of the wall without disturbing the EHT mixture. On the following day, culture media was replaced to maturation medium containing higher Ca²⁺, lipids, low glucose, and supplements to match RPMI composition as described.^20^ Maturation media was based on DMEM low glucose with GlutaMAX™ Supplement and sodium pyruvate (DMEM, Gibco 10567014) containing 1X B27, 1X penicillin/streptomycin, 10% KOSR, 5 µg/mL cobalamin (Thermo Scientific Chemicals A1489403), 0.82 µM biotin (Thermo Scientific Chemicals A1420703), 1X non-essential amino acids (Gibco 11140-50), and 1X Normocin (InvivoGen ant-nr-2). Media were refreshed every 2-3 days. From day 7 to 21, 100 nM triiodothyronine (Millipore Sigma T2877) and 1 µM dexamethasone (Cell Signaling Technologies 14776S) were added.^21^ One-month old EHTs were used for all optical mapping experiments.

### 4.4. Immunofluorescence

EHTs were fixed with methanol at -20 °C for 20 min, followed by blocking and permeabilization in a solution containing 10% normal goat serum (Southern Biotech 0060-01) and 0.5% Triton X-100 in 1X phosphate buffered saline (PBS) for 1 h. Primary antibodies were incubated for 2 h at 30 °C with mouse anti-sarcomeric α-actinin (Invitrogen MAI-22863) and rabbit anti-vimentin (Cell Signaling 5741S) both 1:200 dilution, or rabbit anti-sarcomeric α-actinin (Invitrogen 701914) 1:200 dilution and mouse anti-cardiac troponin T (Abcam ab8295) dilution 1:100. Secondary antibodies anti-mouse Alexa Fluor Plus-488 (Invitrogen A32723), anti-rabbit Alexa Fluor Plus-555 (Invitrogen A32732), anti-rabbit Alexa Fluor Plus-488 (Invitrogen A32731), and anti-mouse Alexa Fluor 633 (Invitrogen A21052) used at dilution 1:500 for 2h at 30 °C. DAPI was used for nuclei staining. After staining, EHTs were treated with the TrueView autofluorescence quenching kit (Vector Laboratories SP-8400-15) according to manufacturer instructions and mounted using Vectashield Vibrance mounting media (Vector Laboratories H-1700-2). Imaging was performed using a Nikon AXR/NSPARC confocal microscope, with a 4X and 60X oil objective.

### 4.5. Contraction assessment

C-Pace and 6-well carbon electrodes (IonOptix) were used to deliver field electrical stimulation at 1, 2, and 3 Hz biphasic 12 V 2 ms pulses. Brightfield recordings were obtained with an inverted microscope equipped with a 4X objective and an IDS U3-3130 CP camera controlled with IDS Python library. Acquisition was performed at 200 fps at 1, 2, and 3 Hz with 60 seconds recording for each step. Pixel size was calibrated using a stage micrometer. Contractility was quantified with BeatProfiler.^22^

### 4.6. Optical mapping perfusion-stage mount

The perfusion stage comprises a base, tissue holder, clamps, and a slider to deliver point electrical stimulation (**Fig. 2C**). The base was printed in two parts including a perfusion bracket to hold a ThermoClamp (AutoMate Scientific) that was epoxied to the bottom part.

The tissue holder was printed in three parts including the detachable tissue holder and two perfusion mounts. Perfusion mounts were used to connect 16-gauge blunt-end needles that were bent and shaped to optimize perfusion. The outlet opening was finished with a rounded 45° bevel to ensure smooth vacuum aspiration. The two inserts were secured with M2X6mm nuts glued to the detachable tissue holder. The detachable tissue holder has a shallow central divot to hold a single milliPillar module containing an EHT, while two clamps prevent flotation during perfusion. A temperature probe was glued next to the tissue divot to monitor local temperature throughout the experiment.

A ThermoClamp perfusion pencil (AutoMate Scientific) was used to warm bathing solution to 37 °C throughout imaging. To maintain temperature during recording, wherein perfusion stops to avoid motion artifacts, we connected a heater under the detachable tissue holder. A flat piece of scrap metal was glued in place and a heating element with a feedback temperature probe were attached (AutoMate Scientific). The inner chamber was then encapsulated in back electronics epoxy (MG Chemicals, 832B). Both pencil and stage heater were controlled with a dual ThermoClamp 3.2 heating system (AutoMate Scientific). A setpoint of 45 °C for both channels was sufficient to maintain 37 °C within the EHT. Throughout all experiments, the temperature of the encapsulated heater remained below 45 °C, and no deformation of the detachable tissue holder was observed. Nevertheless, users should independently verify the thermal behavior of their own setup to ensure that no overheating occurs under their specific operating conditions. The mount was designed in Tinkercad (Autodesk) and printed using an Ender 3 V2 Neo 3D connected to a filament dryer box containing black polyethylene terephthalate glycol (PETG) 1.75 mm filament (Creality). STL files were converted to G-code using UltiMaker Cura and printed at a layer height of 0.12 mm, 100% infill for tissue holder and clamps and 20% for base, 240 °C for nozzle and 70 °C for base, at a speed of 50 mm/s.

### 4.7. Solutions and dye loading

Tyrode’s solution (135 mM NaCl, 5 mM KCl, 1.8 mM CaCl_2_, 1 mM MgCl_2_, 5.6 mM glucose, and 10 mM HEPES, buffered to pH 7.4 with 10 M NaOH) was employed for rinsing, loading and perfusion. Blebbistatin (5 µM, Selleckchem S7099) was added to suppress contractions during optical mapping. The dual-dye loading cocktail contained 30 µM RH-237 (Invitrogen S1109), 10 µM Rhod-2 AM (Invitrogen R1244), and 0.1 % Pluronic F127 (Invitrogen P6867). For pharmacological validation, 100 nM E4031 (MedChemExpress HY-15551) was added to Tyrode’s solution containing blebbistatin.

To model VT, we used a low K⁺/Mg²⁺ modified Tyrode’s solution (135 mM NaCl, 2 mM KCl, 1.8 mM CaCl_2_, 0.5 mM MgCl_2_, 5.6 mM glucose, and 10 mM HEPES, buffered to pH 7.4 with 10 M NaOH) plus 500 nM E-4031. All reagents, except E4031, were dissolved in dimethylsulfoxide (DMSO, Millipore Sigma D2650). EHTs were individually loaded in 1 mL loading solution in a sealed micro-centrifuge vial at 37 °C for 20 min, rinsing with fresh Tyrode’s Solution before and after loading.

### 4.8. Optical mapping imaging protocol

Loaded EHTs were positioned in the central divot, secured with EHT holders, and perfused at 37 °C. Biphasic 2 ms pulses (4 V) were delivered via point electrodes overlapping one half of a pillar head. EHTs were paced from 1 to 3 Hz in 0.5-Hz increments. For each frequency, perfusion continued for 45 s followed by a 15 s pause to minimize fluid-induced motion. Dual-channel voltage and Ca²⁺ images were acquired simultaneously at 1000 fps for 7 s using a SciMedia MiCAM03 system with the lowest illumination intensity that provided adequate focus. After each recording, perfusion resumed and tissue response was confirmed in SciMedia BV Workbench.

For imaging, a TTL generator programmed on Arduino microcontroller triggered pacing for 1 min and during acquisition. We programmed a keypad to have protocols at different pacing frequencies for quicker experimentation. The microcontroller also controlled LED illumination and camera operation, while halting perfusion to avoid motion artifacts. An Arduino case was obtained from The Optogenetics and Neural Engineering (University of Colorado) website.^51^

### 4.9. Optical mapping data analysis

We carried out all data processing, analysis, and figure generation in Python using the following libraries: NumPy, SciPy, OpenCV, scikit-image, Pillow, pybaselines, and matplotlib.

#### Denoising, baseline correction and normalization

Baseline electrophysiology (7 s) were processed sequentially with gaussian spatial denoising (σ = 2), third-order Butterworth temporal band-pass filtering (0.1-25 Hz) and a 3-frame temporal median filter within a binary tissue mask. Baseline drift was removed by Asymmetric Least Squares (ASLS, p = 0.01), with a λ scaled inversely with frequency (5·10^8^ at 1 Hz to 5·10^7^ at 3 Hz). After subtraction, a third-order Savitzky-Golay filter smoothed the signal using an adaptive window (51-15 ms for 1-3 Hz). Signals were normalized using a percentile-based approach (1st to 99th) to minimize the impact of pixel outliers.

For CTL and VT recordings (19 s) a more aggressive pipeline was used: gaussian spatial denoising (σ = 4), band-pass filtering (0-25 Hz), 5-frame temporal median filter, ASLS (p = 0.01, λ=2.5·10^6^), and Savitzky-Golay (9 ms). Parameters were chosen to preserve EAD morphology. Signals were normalized using 0-100 percentiles and the normalized stack was saved for downstream analysis. ASLS baseline suppression effectively eliminated signal drift and diastolic floor shifts associated with increasing beating frequency of the VT condition. Peak amplitude decreased over time due to progressive photobleaching in long recordings or increases in beating frequency observed in VT condition. Per-pixel renormalization was applied in areas with higher beating frequencies using sufficient padding to avoid edge-related artifacts.

#### Parameter calculations in baseline electrophysiology recordings

Four equally spaced 7 × 7 pixels ROIs were placed along the longitudinal axis of each EHT and used to measure APD_80_, CaD_80_, tau, and CV. Resulting trace for each ROI was smoothed (Savitzky-Golay window = 21 frames, order = 3). Peaks were detected on the smoothed trace using ”SciPy find peaks” with a height >0.3X median intensity, prominence ≥ 0.2, and minimum inter-peak distance of 150 ms; peaks were further pruned by valley depth < 40 % of the preceding peak and a relative height ≥ 40 % of that peak, followed by a hard 200 ms binning that retained only the tallest peak per window. Beats were assigned from ROIs that aligned within ± 200 ms with propagation direction starting from point stimulation (right side, ROI1). For each accepted beat: peak time, dV/dt_max_, and D80 (interval between 20% peak and 80% return to baseline) for voltage recordings with additional tau and R^2^ from a mono-exponential fit to the 20-80% segment for Ca²⁺ recordings. Pairwise ROI1-4 were converted to cm using a pixel size of 0.0445 µm/pixel and used to calculate velocities based on activation times calculated by dV/dt_max_ time between extreme ROIs. All beat-level metrics from each ROI per beat were filtered using an interquartile-range outlier filter (k=3) and summary statistics were calculated.

#### Ensemble activation and APD_80_ and DF maps for baseline electrophysiology recordings

Pixel-wise activation times were obtained from the first derivative peak of each beat after frame-wise Gaussian filtering (σ = 1). Outliers were removed via IQR, then maps were normalized to the earliest activation, upsampled 4X with cubic interpolation, and median-filtered (9 × 9) for display. Ensemble activation maps were averages of normalized beat-wise maps. APD_80_ maps used a uniform box blur (size = 9) followed by 3 × 3 median filtering; APD_80_ was defined as the interval between 20 % upstroke and 80 % repolarization per pixel, with IQR outlier removal and beat-wise averaging. Dominant frequency (DF) maps were generated after Gaussian spatial filtering (σ = 2) and Savitzky-Golay temporal smoothing (51 ms, order 3); DF was the frequency of maximal FFT amplitude within 0.2-6 Hz for each masked pixel. Ensemble DF statistics and cumulative spectra were exported as CSV; DF maps were median-filtered (9 × 9) only for visualization.

#### Measurements of AP-CaT latency in baseline electrophysiology recordings

KairoSight^52^ was used to determine AP-CaT latency on the first beat of recordings at a 500 ms CL (**Fig. 2J-M**).

#### Parameter quantifications in Kymograph in CTL and VT recordings

Median fluorescence along a 3-pixel line spanning the longitudinal axis generated voltage and Ca²⁺ fluorescence time-distance maps. Kymographs were displayed using a robust normalization (rolling 4 s) to mitigate drift. Raw traces were used for calculations after a 7 ms temporal gaussian temporal smoothing for APD and DI and 9 ms for dV/dt. Beat-wise activation times were defined by dV/dt_max_ per distance column. CV was derived from linear fits of activation time versus distance. APD_80_ used a 20% upstroke to 80% repolarization threshold, and diastolic interval (DI) was the interval from the prior-beat’s 80% repolarization to the next upstroke onset.

#### Beat-resolved activation, APD, dV/dt maps and DF in CTL and VT recordings

Voltage signals were smoothed (Savitzky-Golay, 9 ms window, order 3). Activation time per pixel was calculated from maximum dV/dt times. To ensure that activation, repolarization, and upstroke metrics were extracted from the same physiological event across the field of view, a ROI-guided wave-tracking procedure was applied. ROI-guided wave tracking selected a single propagating wave: activation candidates from ROI traces defined expected arrival times, and the activation of each pixel was taken as the local dV/dt maximum within ±120 ms (±140 ms for APD). Activation times were reported in milliseconds and referenced to the earliest activated pixel (time zero). APD_80_ was defined as the time between 20% upstroke and 80% repolarization. The dV/dt max map reported the upstroke velocity magnitude. Outliers were removed with IQR (k = 1.5) and gaps filled; spatial blurring (σ = 1) was applied for display. DF maps were computed per masked pixel using Welch PSD (Hann window, ≈2 s segments, 50% overlap) after Savitzky-Golay smoothing (7 ms). The dominant frequency was the peak within 0.5-20 Hz; post-processing used IQR (k = 1.5) and σ = 1 blurring. Maps were exported as overlays and per-pixel tables.

#### Phase singularities (PS) and wavelets in CTL and VT recordings

For Hilbert-based phase analysis, tissue mask was eroded by 1 px to reduce boundary artifacts, and the image stack was spatially smoothed using an edge-safe masked Gaussian filter (σ = 4), computed as Gaussian (signal × mask) / Gaussian(mask) to avoid blurring across the tissue boundary. For phase calculations, each pixel trace was detrended by subtracting a 0.8 s running-mean baseline and mean-centered. Pixels with low variability were excluded using an adaptative SD threshold (SD ≥ max (10^-3^, 0.25X median SD within tissue)), and retained signals were standardized by their SD. Instantaneous phase (φ) was obtained from the analytic signal computed by a temporal Hilbert transform and defined as the angle of the analytic signal. PS were detected per frame using a winding-number method on 2×2 pixel cells fully contained in the valid mask by summing wrapped phase differences around each cell; candidates with |ΣΔφ| < 1.8π were rejected, and remaining detections were assigned topological charge of either clockwise (+1) or counter-clockwise (−1) based on the direction of the phase wrap. To reduce grid bias, PS detection was performed on a half-pixel-shifted lattice (phase was reconstructed after bilinear interpolation of cos(φ) and sin(φ) with a + 0.5 px shift in x and y to avoid wrap artifacts), and only regions with near-complete mask support after shifting were evaluated. Single PS per connected component (≥ 1 px) was placed at the centroid; detections within 5 px were merged via dilation and connected-component clustering.

PS tracks were built by greedy nearest-neighbor association respecting chirality, allowing ≤ 5 px inter-frame displacement, gaps up to 10 frames, and a minimum lifetime of 100 ms. Per-track summary data was calculated containing duration, cumulative path length (sum of stepwise Euclidean displacements between successive observed positions), net displacement (start-to-end distance) and mean speed (path length divided by duration). Pixel distances were converted to physical units using the experiment-specific spatial calibration (0.0445 px/µm), with distances optionally reported in µm and speeds in derived physical units. Tissue-level summary statistics were obtained by pooling tracks within each tissue and reporting pooled medians/maxima of duration and motion metrics, rotor incidence rates normalized by total recording duration (rotors per minute), and peak rotor concurrency based on overlapping track intervals.

#### Anatomy normalization and cohort-level dwell time and prevalence mapping

Each tissue mask was aligned by PCA: centering at the centroid, rotating to align the first principal axis with the long axis, and mirroring to enforce consistent left-right orientation. PS coordinates were transformed into this frame and expressed in µm (0.0445 px/µm). Per-tissue dwell maps binned anatomy-normalized PS positions (10 µm bins); bin counts were multiplied by Δt = 1/FPS (ms) for plotting. Mean dwell maps summed per-tissue histograms and divided by the number of tissues with rotors (equal weighting). Prevalence maps marked bins visited at least once per tissue and calculated the percent of tissues showing occupancy. Both maps were rendered within a canonical EHT silhouette that was derived from the ensemble of aligned, length-normalized tissue masks.

Wavelet (wavefront component) quantification and PS-wavelet overlays: Normalized voltage recordings were renormalized using the 2nd-98th percentiles of masked pixels, and Gaussian-smoothed (σ = 4) to remove outlier and further reduce noise. Instantaneous phase was obtained using temporal Hilbert transform. At each frame, a wavefront mask comprised pixels with phase distance to 0 ≤ 0.40 rad within the tissue mask and valid phase. Wavelets were defined as contiguous blobs of wavefront pixels, where pixels were considered connected if they touched by an edge or a corner (8-connectivity) with a minimum area of ≥ 25 px. To avoid over-fragmentation, only components whose area was at least 25% of the largest component in that frame were retained. Wavelet counts per frame were obtained from the filtered set. Wavelet tracks were built by greedy one-to-one overlap matching: predecessor-successor pairs with intersection/smaller-component area >0.20 were linked. Tracks shorter than 30 ms were discarded; the resulting persistence-filtered wavelet count time series was aligned with PS data for overlay plots and Suppl. videos.

### 4.10. Statistical analysis

All statistical analyses were performed on tissue-level summary values (one value per tissue per condition), and data are presented as boxplots with the center line denoting the median and the box spanning the interquartile range (25^th^-75^th^ percentiles); sample sizes are reported in the figure legends. Normality was assessed using Shapiro-Wilk. When normality assumptions were not met for any subpanel within a figure, all subpanels for that figure were analyzed using nonparametric tests to maintain a consistent statistical approach.

Paired versus unpaired testing was determined by the data structure: experiments with complete within-tissue measurements across levels were analyzed with paired methods, whereas datasets with incomplete pairing (e.g., heterogeneous frequency sets and exclusions for insufficient response) were treated as unpaired.

For paired experiments with >2 groups differences were tested using the Friedman test, followed by paired Wilcoxon signed-rank comparisons versus 1 Hz (**Fig. 1**). Frequency-response optical mapping data used the Kruskal-Wallis test with Dunn’s post-hoc comparison vs 333 ms (**Fig. 2**). In **Fig. S3**, with normal distribution, differences were assessed using a paired two-sided t-test. In **Fig. 3** paired metrics were analyzed using a two-sided paired Wilcoxon signed-rank test because several paired differences deviated from normality. For **Fig. 4**, some data did meet normality assumptions, so a two-sided unpaired Mann-Whitney U test was used. These data were also treated as unpaired because recordings were not available at identical frequency sets for all tissues as discussed in the results. A p < 0.05 was considered statistically significant. No formal correction for multiple comparisons was performed.

## Declaration of generative AI and AI-assisted technologies in the writing process

During the preparation of this work, the author(s) used OpenAI models to improve readability and grammar. After using this tool/service, the author(s) reviewed and edited the content as needed and take full responsibility for the content of the publication.

## Acknowledgments

We thank the Advanced Cellular and Tissue Microscopy Core and Research Pathology Core at Houston Methodist. This work was supported by an American Heart Association (AHA) career development award (19CDA34680003, FA), National Heart, Lung, And Blood Institute of the National Institutes of Health (NHLBI) R01HL158703 and R01HL168277 (FA), Houston Methodist Cornerstone Award (MV, FA), and the Houston Methodist Startup Funds (FA).

## Conflict of Interest

The authors declare no conflicts of interest.

## Data Availability Statement

The data that supports our findings in this study remains available from the corresponding author upon reasonable request.

## SUPPLEMENTAL VIDEO LEGENDS

**Supplemental Video 1**. Handling of milliPillar modules. Video shows how to detach milliPillar modules from multi-well plates using forceps.

**Supplemental Video 2**. Representative contractility trace. *Left panel*, contractility recording video. *Middle panel*, BeatProfiler-generated video showing pillar tracking over time. *Right panel*, show quantification of force overtime. Videos were recorded at 1, 2 and 3 Hz, for 60 s each. Orange segmented line represents the baseline at 1 Hz

**Supplemental Video 3**. Dual voltage and Ca²⁺ optical mapping with point stimulation at 1 Hz (CL = 1000 ms). *Top panel*, Voltage and Ca²⁺ maps showing wave propagation across the EHT. A jet-style color scale (blue to red) depicts normalized signal intensity over time. *Bottom panel*, simultaneous normalized signal quantification at ROI1, 2 and 3. Point electrical stimulation was applied on the right side of the EHT at 1 Hz.

**Supplemental Video 4**. Dual voltage and Ca²⁺ optical mapping with point stimulation at 3 Hz (CL = 333 ms). Same as described for **Supplemental Video 3**.

**Supplemental Video 5**. Representative CTL recording show synchronized capture throughout the entire longitudinal axis due to field stimulation. *Left panel*, Normalized voltage and Ca²⁺ fluorescence (normalized 0-1 in turbo color scale), *Right panel,* phase calculated from these recordings (scale -π to π, in turbo color scale). The video corresponds to the ∼15-19 s interval from the Kymograph on **Fig. 4C** top panel is shown

**Supplemental Video 6**. Events occurring between ∼15-19 s from the representative VT recording shown in **Fig. 4C**, which shows activation originating from the top-left corner of the EHT that progressively slows as the beating frequency accelerates before wavebreak, with partial recovery towards the end of the recording. *Left panel*, Normalized voltage and Ca²⁺ fluorescence (normalized 0-1 in turbo color scale), *Right panel,* phase calculated from these recordings (scale -π to π, in turbo color scale).

**Supplemental Video 7**. Events occurring between ∼10-14 s from the representative VT recording shown in **Fig. 5A**. Video shows from beats 12-19 and the formation of rotos on both EHT heads as described in **Fig. 5I**. *Left panel*, Normalized voltage and Ca²⁺ fluorescence (normalized 0-1 in turbo color scale), *Right panel,* phase calculated from these recordings (scale -π to π, in turbo color scale).

**Supplemental Video 8:** Wavelet and PS localization from Fig. 7 showing events occurring between ∼0-6 s. *Top panel*, voltage signal (normalized 0-1 in turbo color scale), *middle panel*, phase calculations (scale -π to π, in turbo color scale) with wavelet outline overlap in white (kept) or orange (excluded because did not meet size or time as described in Methods), *bottom panel*, binary wavelets mask. PS are shown as circles when they met 1.8π criteria as described in the Methods section.

## SUPPLEMENTAL STL FILES

**STL file 1.** Modified milliPillar mold including rafts and supports.

**STL file 2.** Perfusion stage (part 1/6). Perfusion heater holder for ThermoClamp pencil.

**STL file 3.** Perfusion stage (part 2/6). Base to hold perfusion heater holder and tissue perfusion mount. Use epoxy to glue perfusion heater holder to base.

**STL file 4.** Perfusion stage (part 3/6). Tissue perfusion mount.

**STL file 5.** Perfusion stage (part 4/6). Perfusion holder to mount blunt needles.

**STL file 6.** Perfusion stage (part 5/6). Clamp to hold EHTs.

**STL file 7.** Perfusion stage (part 6/6). Electrical stimulation slider.

**Table 1.**
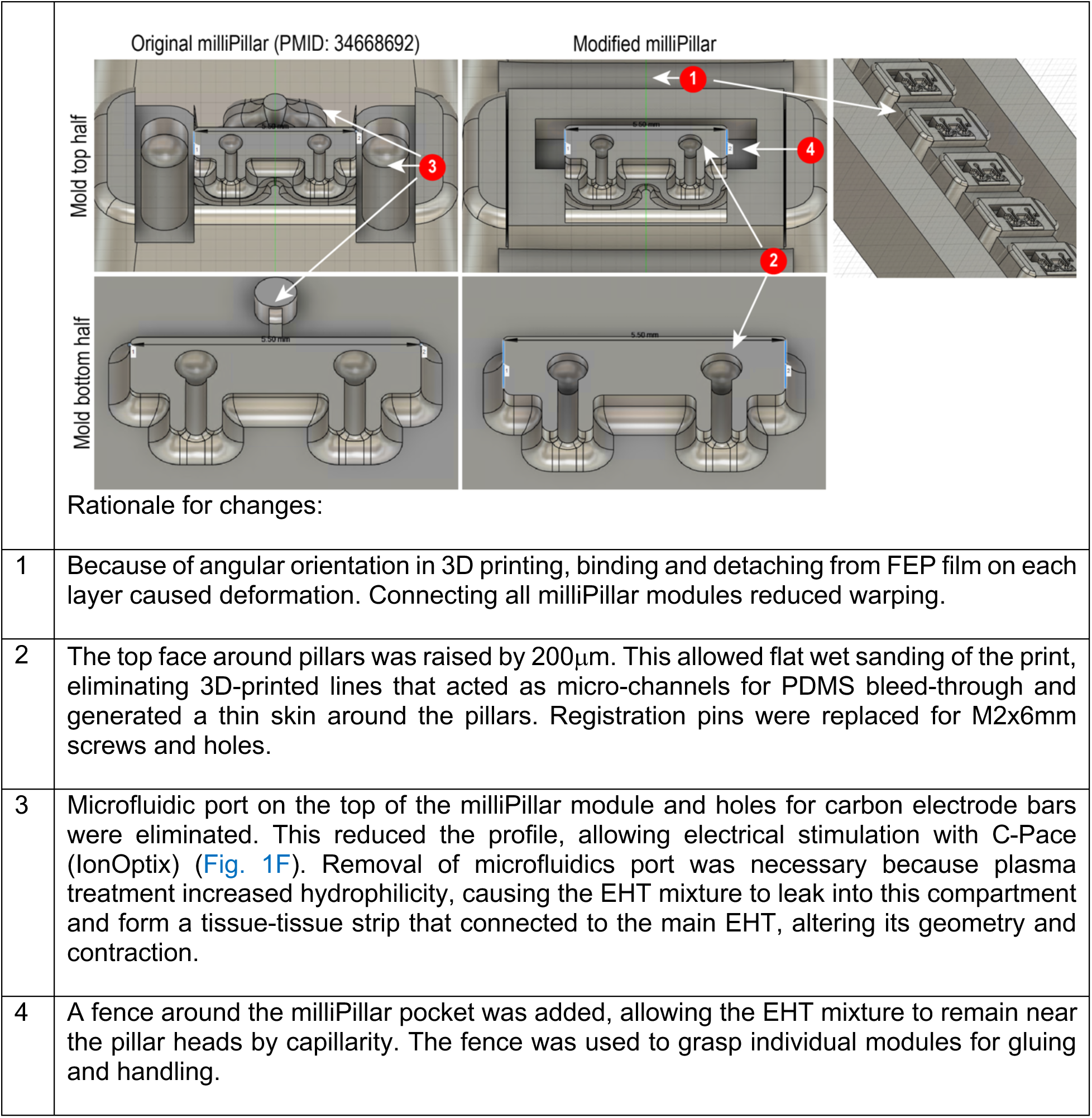
Summary of changes to milliPillar. Original CNC files from milliPillar publication^17^ were slightly modified using Fusion 360 (Autodesk). Images shown below were obtained using Fusion 360.

**Table 2.**
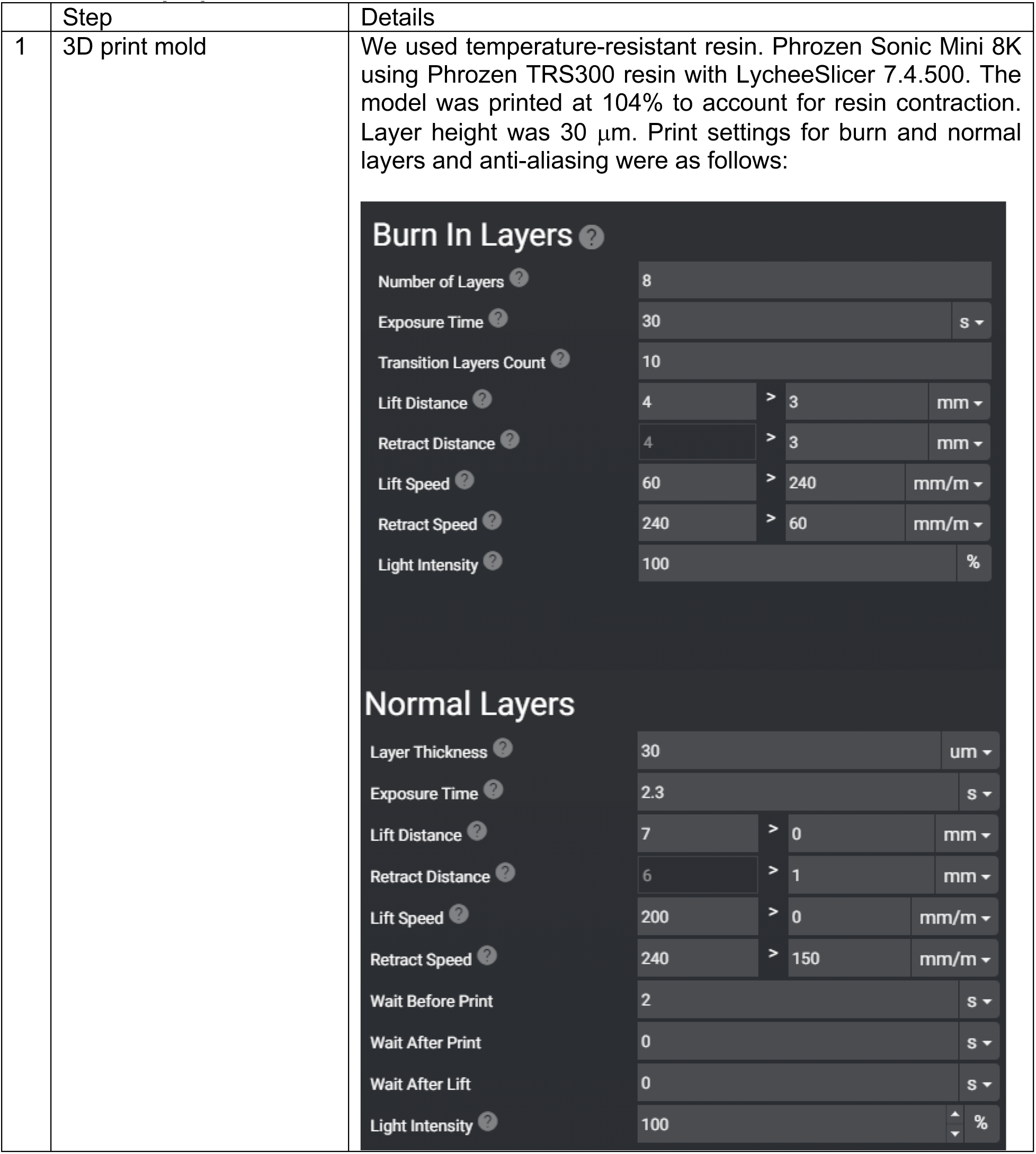
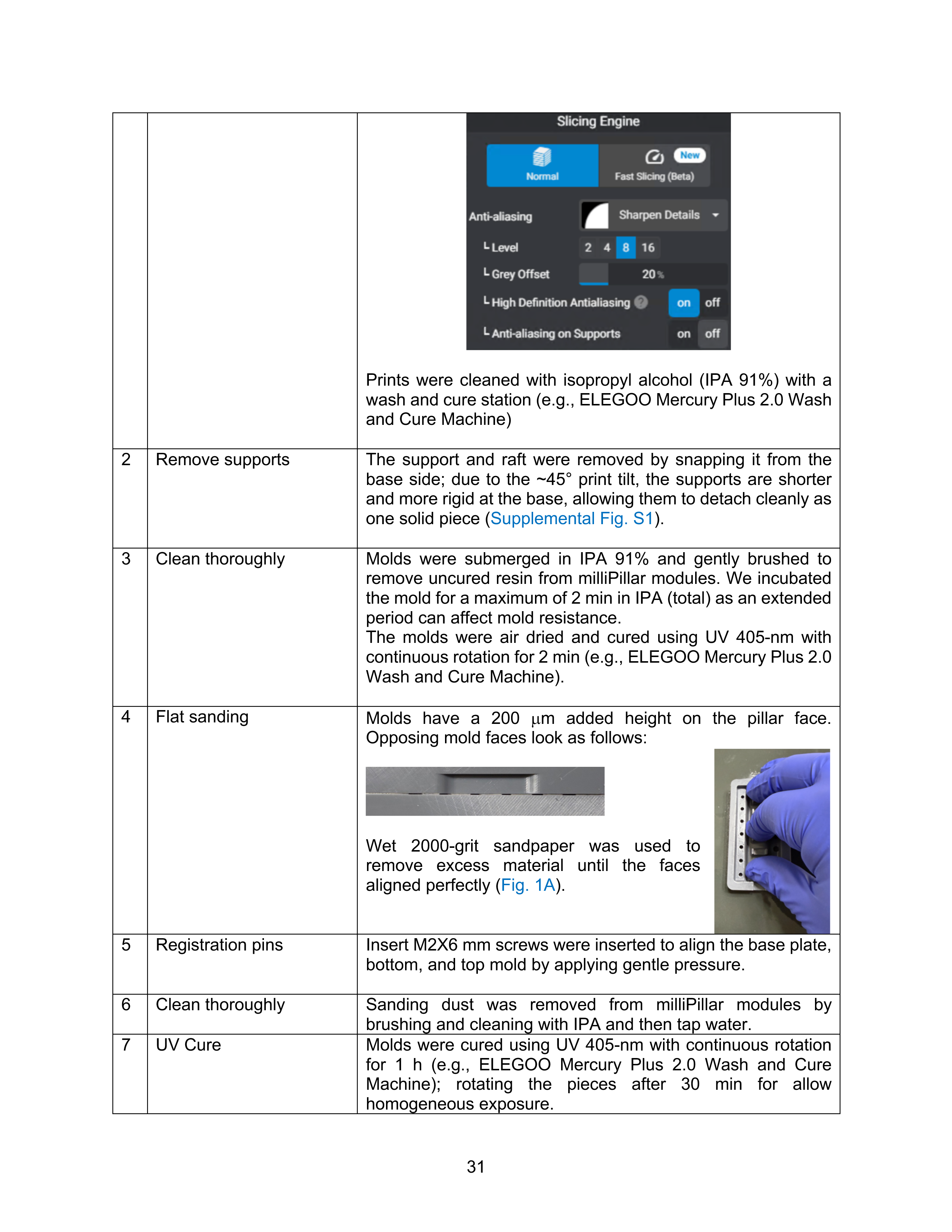
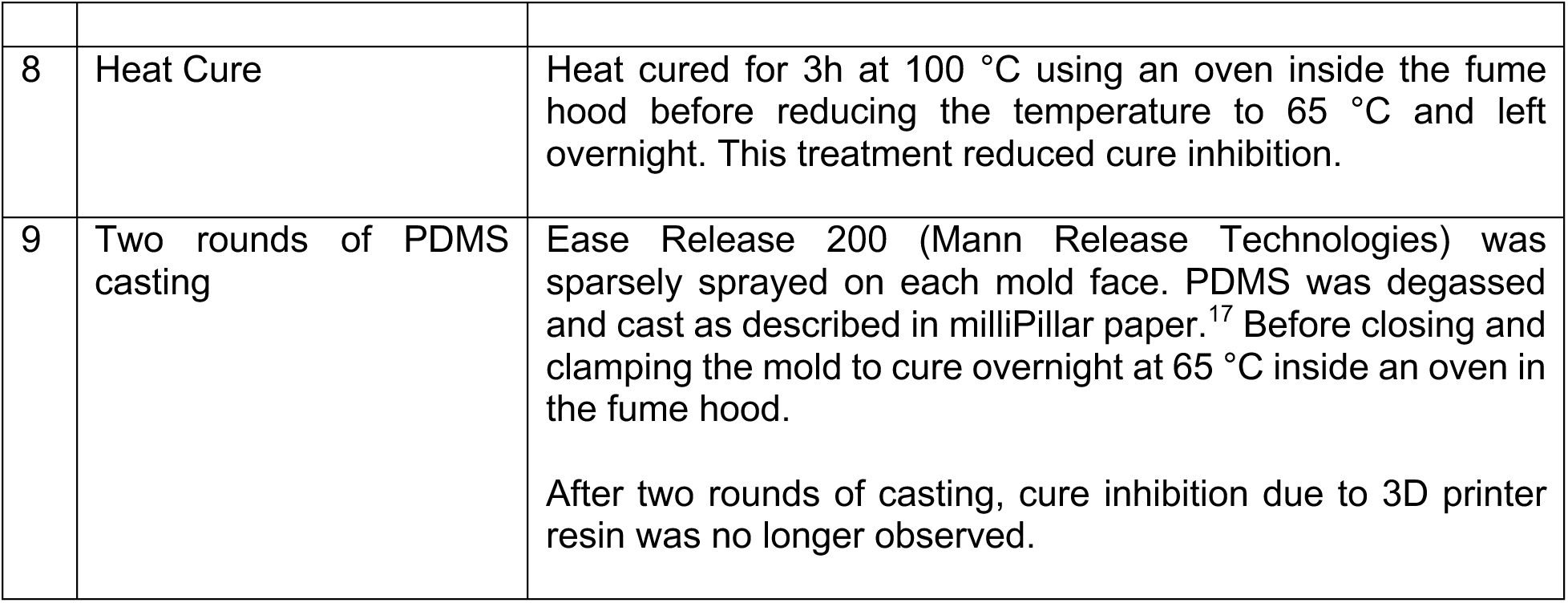
Mold preparation.

**Table 3.** PDMS for EHT preparation.

|  | Step | Details |
| --- | --- | --- |
| 1 | Clean mold thoroughly and apply release agent | Ease Release 200 (Mann Release Technologies) was sparsely sprayed on each mold face to help unmold PDMS. |
| 2 | Cast PDMS | PDMS was prepared with base-to-curing-agent 10:1 (Sylgard™ 184, #24236-10, Electron Microscopy Sciences), degassed, poured into the molds and degassed again. |
| 3 | Clamp and cure | Follow the milliPillar publication for details.<sup>17</sup> Briefly, molds were clamped using C-clamps (#1705A11, McMaster) and a 1/4" × 4" hex standoff (#91780A060, McMaster). Molds were topped off with PDMS and incubated at 65 °C overnight. |
| 4 | Unmold | Because of modifications, the PDMS platform is more prone to break compared to original milliPillar. Molds were gently removed by separating the sides of the platform with plastic pipet tips. The PDMS platform was removed and then further cured at 65 °C for additional 24 h. |
| 5 | Cleaning | After curing, molds were washed with MilliQ water, 70% ethanol and MilliQ water, then dried at 65 °C. |
| 6 | Cut and glue | Using a scalpel, modified milliPillar modules were cut and kept aligned on a clean and dust-free surface (Fig. 1A, Supplemental Fig. 2). Using a sterile L-shape cell spreader, a few drops of PDMS were spread on a clean sterile 6-well plate lid to make a thin film that was barely wet (Supplemental Fig. 2A). Then, forceps were used to grab the modified milliPillar fence (Supplemental Fig. 2B) and wet the bottom of each module with PDMS (without affecting pillars) and then transferred each module into a 6-well plate.<br>Spreading the thin layer of PDMS was the most effective and simple way to avoid excess PDMS wetting and filling between plate and milliPillar module. Supplemental Fig. S2C and S2D show examples of incomplete filling and overfilling. Excess PDMS interfered with EHT plating, as these thin films detach when plating and get incorporated to the tissue or added extra material that deformed pillar heads (Supplemental Fig. S2C). Incomplete filling introduces an additional issue, as plasma treatment increases surface hydrophilicity, making the mixture prone to leaking underneath the milliPillar module.<br>To prevent plasticware deformation, the thin layer was cured overnight at 45 °C. |
| 7 | Corona plasma treatment | Corona plasma was delivered inside and around the milliPillar module using a spring tip wire, and a Laboratory Corona Treater (#12201 and #BD-20AC, Electro-Technic Products). |
| 8 | Wash and decontamination | Wells were washed twice with 70% ethanol. Ethanol was incubated for 30 minutes inside the biosafety cabinet and then aspirated and air-dried.<br>Plates and lids were treated for 20 min with UV-C in a UV Clave (B1450, Benchmark Scientific; 600mJ/cm<sup>2</sup>, 20 min cycle) inside the biosafety cabinet. UV-C indicator stickers (B1450-ISS, Benchmark Scientific) monitored UV-C efficiency and penetrance. UV-C went under milliPillar PDMS but did not cross the 6-well plastic lid. Therefore, both plate and lid were treated separately, with inner faces exposed directly to UV-C. |
| 9 | Blocking | KOSR blocking prevented collagen hydrogel sticking to milliPillar module, removing the need to needle-score each individual EHT at 1 and 24h post cure as originally described.<sup>17</sup> This made plating faster and EHT formation more reproducible.<br>A volume of 25 µL of KOSR was enough to coat the entire milliPillar pocket. After 1 h incubation at 37 °C, we aspirated and dried KOSR. It is important that no leakage is observed at this step between the interface of PDMS and plastic. Drying well KOSR before plating prevents cell-hydrogel mixture leak underneath by capillarity. However, it is recommended to properly glue milliPillar inserts as described above and in Supplemental Fig. S2. |
| 10 | Plating | Confluent hiPSC-CMs and cardiac fibroblasts were detached using TryPLE Select 10x supplemented with 5 µM Y-27632 for approximately 15 min. Equal volume of plating media (RPMI, 1x B27, 10% KOSR, 1X Pen Strep, 5 µM Y-27632) was added and pipetted up and down, spun and resuspended in 5-10 mL of fresh plating media.<br>A total of 550,000 cells (75% cardiomyocytes/25% fibroblasts) were used for each EHT. Batches of at least 12 EHTs were plated simultaneously. For each EHT, cells were pelleted and resuspended in 15 µL of matrix solution per EHT, consisting of 12 µL of PureCol® EZ Gel (BICO 5074, 5 mg/mL bovine atelocollagen) and 3 µL plain RPMI supplemented with 25 µM Y-27632, to achieve a final collagen concentration of 4 mg/mL and 5 µM Y-27632.<br>EHT solution (15 µL) was homogeneously dispersed without introducing bubbles and applied to each milliPillar module pocket near the pillar heads by sweeping the pipette tip across the pocket and dispensing on the opposite pillar. We also spread the mixture around the upper fence to ensure the |
|  |  | mixture remained around pillar heads by capillarity. A 25 μL of plating media was added around the plastic well borders to add extra moisture and prevent dehydration.<br>Cell-collagen hydrogel mixture was incubated for 1h at 37 $^{\circ}$C to allow collagen polymerization and then 4 mL of fresh plating media was added against the wall to avoid disturbing the mixture. |
| 11 | Maturation | After 24 h of plating, media was replaced with maturation media based on DMEM low glucose containing 1X B27, 1X penicillin/streptomycin, 10% KOSR, 5 μg/mL cobalamin, 0.82 μM biotin, and 1X non-essential amino acids, 1X normocin. Media was refreshed every 2-3 days. From day 7 to 21, 100 nM triiodothyronine and 1 μM dexamethasone were added.<sup>21</sup> One-month old EHTs were used for all optical mapping experiments. |

## SUPPLEMENTAL FIGURE LEGENDS

**Supplemental Figure 1.**
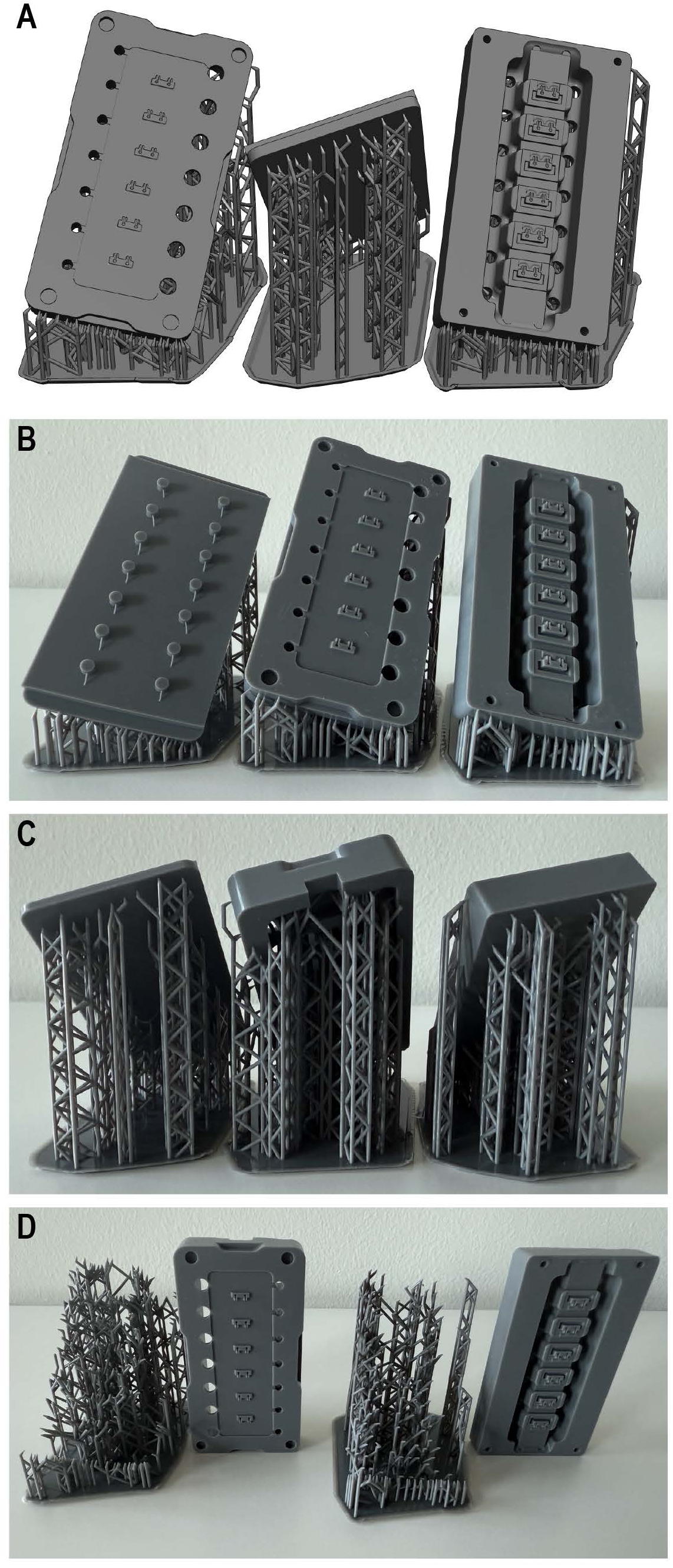
Modified milliPillar platform for resin 3D printing. **A**. 3D rendering of the STL file model showing raft, supports and modified milliPillar. Front (**B**) and back (**C**) views of the printed mold produced with a Phrozen Sonic Mini 8K using temperature-resistant resin (TRS300C, Phrozen). **D.** Raft and supports were easily removed from the modified molds.

**Supplemental Figure 2.**
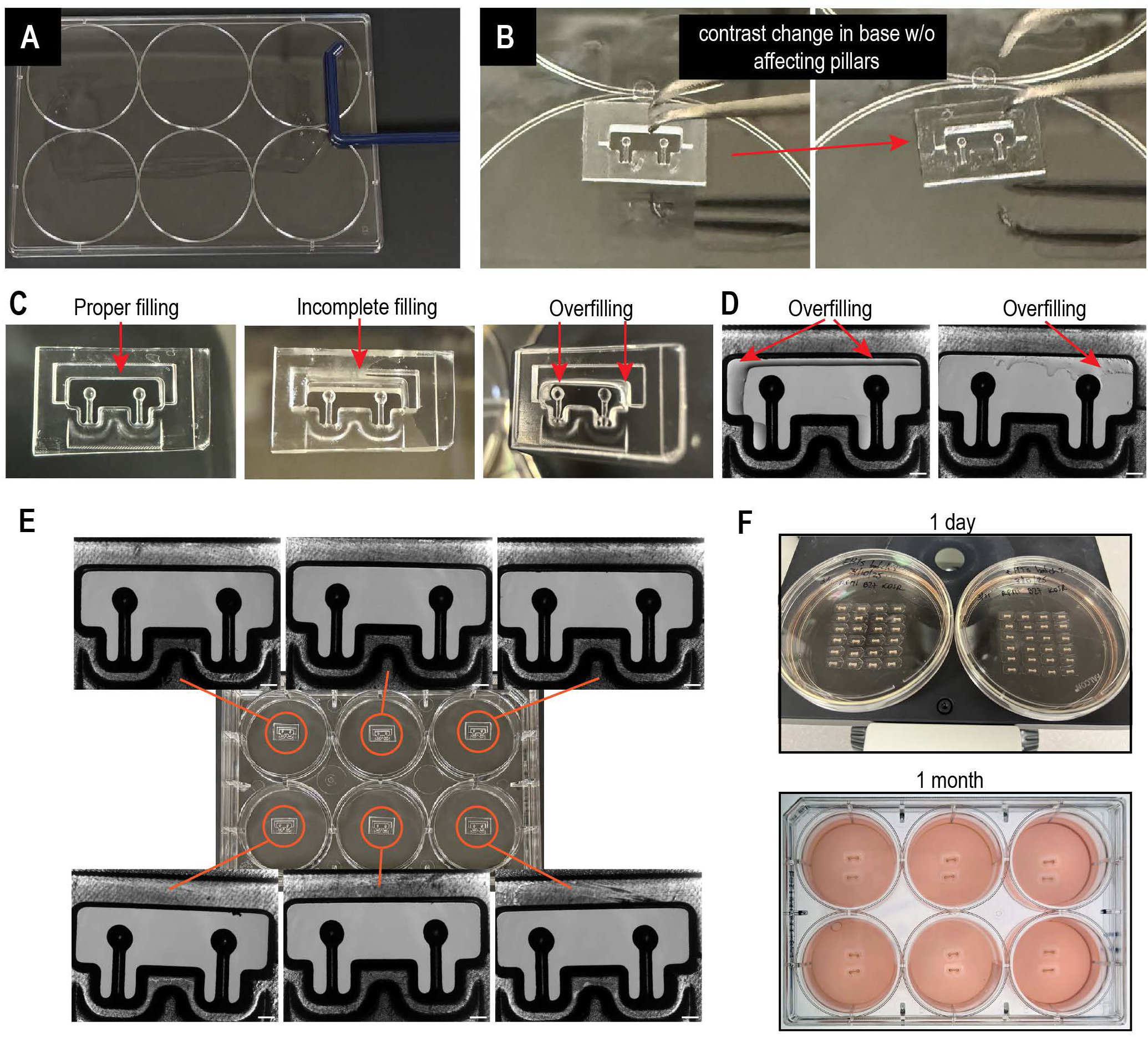
Increased flexibility for the milliPillar platform using individual modules. **A.** A thin PDMS layer was spread on a 6-well plate lid with a L-shaped cell spreader. **B.** Individual milliPillar modules were trimmed, handled with forceps and placed onto the PDMS film for ∼1 s; a subtle contrast change was visible initially at the module base indicating wetting without affecting pillar heads. Proper, incomplete and overfilled interfaces between the module bottom and the 6-well plate are shown (**C**-**D**). **E**. Upside-down view of the 6-well plate and corresponding microscope close-ups demonstrate reproducible attachment across wells. **F**. Using modules as individual units increases versatility, allowing plating from 100 mm dishes to multi-well plates; the subfigure shows complete self-assembly of 40 tissues at 24h after corona plasma/KOSR treatment without the need of needle scoring. Approximately 79% of tissues maintained shape and remained contracting and viable for at least 3 months. Scale bar in microscope images represent 500 μm.

**Supplemental Figure 3:**
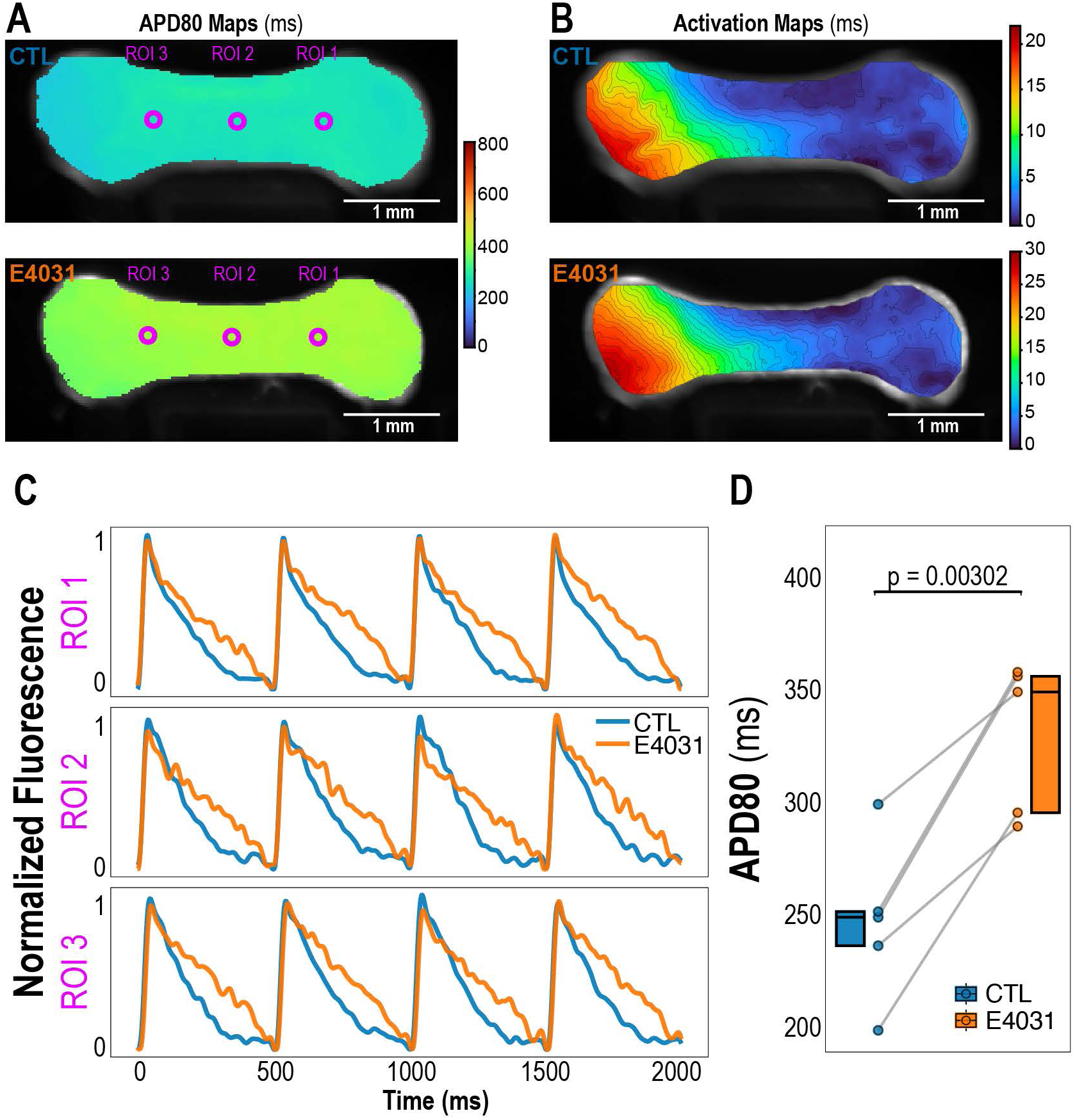
E4031 prolongs APD in EHTs. **A.** Ensemble APD_80_ maps for a representative EHT paced at 2 Hz recorded before (control, top) and after treatment with 100 nM E4031 (bottom). E4031 induced a homogenous APD_80_ prolongation throughout the tissue. **B.** Corresponding ensemble activation maps for the same tissue before (top) and after (bottom) E4031 treatment; note the scale difference between panels. **C.** Normalized voltage traces at each ROI before (blue) and after (orange) E4031 treatment reveals prolonged APD_80_. **D.** Quantification of APD_80_ before and after E4031 treatment. Box plots display the median and IQR with overlaid scatter points representing each individual EHT. N = 5 independent EHTs (across 2 batches). Data were normally distributed, so a paired, two-sided t-test was used.

## Notes

### Competing Interest Statement

The authors have declared no competing interest.

